# Genomic offset is not predictive of recent demographic trends in *Lycaeides* butterflies

**DOI:** 10.64898/2026.06.21.733565

**Authors:** Gbolahan A. Reis, Matthew L. Forister, Lauren K. Lucas, Arthur M. Shapiro, James A. Fordyce, Chris C. Nice, Zachariah Gompert

**Affiliations:** Department of Biology, Utah State University, Logan, Utah, USA; Ecology Center, Utah State University, Logan, Utah, USA; Program in Ecology, Evolution and Conservation Biology, Department of Biology, University of Nevada, Reno, Nevada, USA; Center for Population Biology, University of California, Davis, California, USA; Department of Ecology and Evolutionary Biology, University of Tennessee, Knoxville, Tennessee, USA; Department of Biology, Texas State University, San Marcos, Texas, USA

**Author notes:** **Corresponding Author**: Gbolahan A. Reis, Utah State University, Logan, Utah, USA. **Funding Information**: National Science Foundation, Grant/Award Numbers: DEB-1844941, DEB-2114793, DEB-2114794, Society for the Study of Evolution R. C. Lewontin Early Award, Utah State University Ecology Center Research Award and American Philosophical Society Lewis and Clark fund for exploration and field research.

**Keywords:** Genomic offset, Climate maladaptation, Butterflies, Genotype-environment associations, Climate change

## Abstract

Genomic offset (GO) is increasingly used to predict population maladaptation risk under climate change, with larger offsets assumed to indicate greater vulnerability. Despite rapid adoption in conservation planning, it remains unclear how sensitive GO estimates are to key methodological choices, including SNP set composition, genotype–environment association (GEA) methods, and the specific GO metric used. Empirical validation against observed population dynamics also remains limited. Here, we evaluate the methodological robustness and predictive performance of GO using multidecadal demographic monitoring data from *Lycaeides* butterflies, a system with short generation times and high fecundity that may facilitate rapid adaptive responses. GO estimates were broadly consistent across SNP sets, regardless of composition or size, with climate-associated and randomly selected SNPs yielding largely concordant values. Consistency across GEA methods was moderate and depended on the SNP set used. In contrast, GO metrics differed substantially in the magnitude of maladaptation estimated, suggesting they capture distinct biological signals and should not be treated as interchangeable. Crucially, GO was a poor predictor of observed population trends, regardless of SNP set composition, GO metric, or GEA method, both at sites used to fit GEA models and when extrapolated to independent demographic sites. These findings suggest that, while GO provides a valuable conceptual framework for assessing potential maladaptation, its quantitative estimates and predictive power are sensitive to methodological choices and species-specific biological context. We therefore urge careful alignment of GO metric assumptions with conservation objectives, along with rigorous empirical validation, before GO estimates are used to inform management decisions.

## Introduction

Rapid environmental change, particularly climate change, poses a major threat to global biodiversity by exposing populations to conditions for which they are poorly adapted. In the absence of sufficiently rapid evolutionary responses, such exposure can result in maladaptation and population decline (Aitken *et al*., 2008; Bradshaw & Holzapfel, 2006; Brady *et al*., 2019; Hoffmann & Sgrò, 2011; Nadeau *et al*., 2017; Urban, 2015). The concept of genomic offset (GO) was developed to quantify this risk. Genomic offset is a forecasting tool that predicts the risk of maladaptation by using genotype–environment association (GEA) models to estimate the genomic change required for a population to become well adapted to new environmental conditions. GO can be estimated using various GEA methods. Univariate approaches, such as linear regression and latent factor mixed models (LFMM), test loci independently for associations with environmental predictors. In contrast, multivariate approaches, including redundancy analysis (RDA), gradient forest (GF) and generalized dissimilarity modelling (GDM), consider multilocus associations and can capture covariance among loci and environmental variables (Capblancq & Forester, 2021; Caye *et al*., 2019; Fitzpatrick & Keller, 2015; Frichot *et al*., 2013; Gautier, 2015).

Populations with higher GO values are expected to experience greater reductions in fitness as climate change alters their local environments (Fitzpatrick & Keller, 2015; Gain *et al*., 2023). GO has been applied across continents for diverse taxa including amphibians, birds, mammals, marine invertebrates, crops, and forest trees, and is increasingly recommended for climate adaptive species management (Adam *et al*., 2022; Aguirre-Liguori *et al*., 2019; Bay *et al*., 2018; Bonnier *et al*., 2025; Borrell *et al*., 2020; Bourret *et al*., 2024; Capblancq *et al*., 2020b; Capblancq & Forester, 2021; Chen *et al*., 2022; Dauphin *et al*., 2021; Fitzpatrick & Keller, 2015; Fitzpatrick *et al*., 2021; Francisco *et al*., 2025; Gain *et al*., 2023; Gougherty *et al*., 2021; Gugger *et al*., 2018; Hoste *et al*., 2024; Ingvarsson & Bernhardsson, 2020; Jia *et al*., 2020; Lachmuth *et al*., 2023, 2024; Lind *et al*., 2024; Lind & Lotterhos, 2025; Martins *et al*., 2018; McLennan *et al*., 2025; Rhońe *et al*., 2020; Ruegg *et al*., 2018; Sang *et al*., 2022; Smith *et al*., 2021; Theraroz *et al*., 2024; Vanhove *et al*., 2021; Wu *et al*., 2025). Despite its growing application, the extent to which GO accurately predicts maladaptation risk in wild populations remains uncertain, owing to a lack of empirical validation.

Two conceptually distinct GO metrics are widely used in the literature (Figure 1a, b). The first, originally proposed by Fitzpatrick & Keller (2015), estimates the genomic change required to maintain GEAs when a population transitions from its baseline environment to a new environment. This metric assumes that populations are locally adapted to their baseline environments and therefore that any environmental deviation produces maladaptation. It is calculated as the absolute difference between the expected genomic composition under baseline and projected environmental conditions, where expectations are derived from GEAs estimated in the baseline environment to which populations are assumed to be locally adapted. Importantly, unlike raw environmental distance, which measures the magnitude of change in environmental variables without reference to genetic data, this metric weights environmental variables by their estimated genetic effect sizes before distance is calculated. As a result, environmental changes that are more strongly associated with genomic variation contribute disproportionately to the estimated offset.

**Figure 1:**
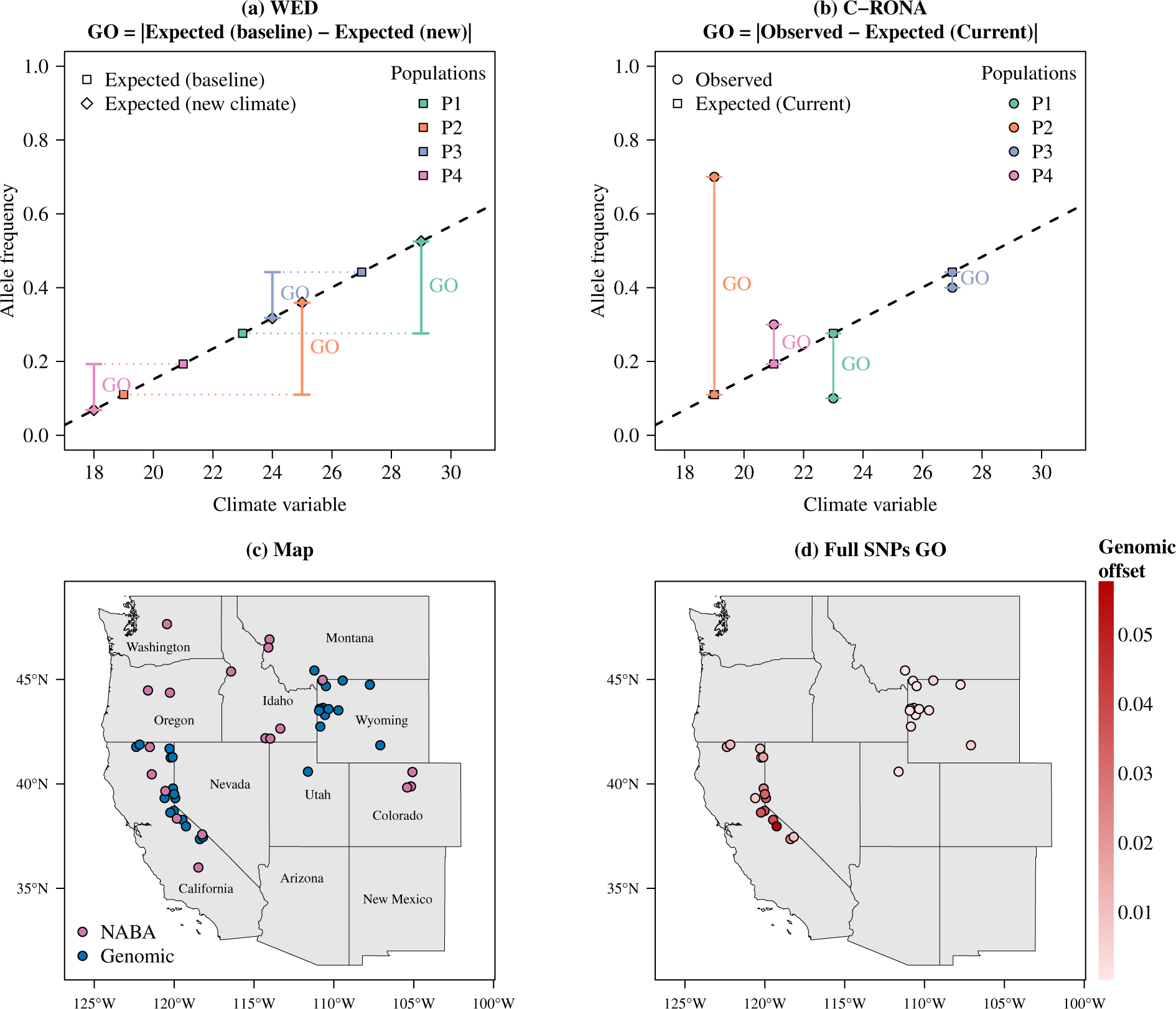
Conceptual illustration of genomic offset (GO) metrics, study area, and example GO estimates. (a) Weighted environmental distance (WED; Fitzpatrick & Keller (2015)) quantifies the genomic change required to maintain genotype–environment associations (GEAs) when a population transitions from its baseline climate, to which it is assumed to be locally adapted, to a novel climate. Unlike simple climatic distance, which measures the magnitude of change in climatic variables without reference to genetic data, WED weights climatic predictors by their estimated genetic effect sizes before calculating distance. Consequently, climatic changes more strongly associated with genomic variation contribute disproportionately to the estimated offset. (b) Current risk of nonadaptedness (C-RONA; Borrell *et al*. (2020)) quantifies the mean absolute deviation between a population’s observed genomic composition and the expected genomic composition derived from GEAs for the target climatic condition. (c) Map of the study area across the western United States showing populations sampled for genomic analysis and long-term NABA monitoring sites from the North American Butterfly Association (NABA). (d) Example WED estimates for each population sampled for genomic analysis, estimated using generalized dissimilarity modeling (GDM) based on the full SNP dataset. Darker red indicates higher genomic offset.

Because this metric effectively represents an genetically-weighted distance in environmental space, we hereafter refer to it as weighted environmental distance (WED). The second metric, termed the current risk of nonadaptedness (C-RONA; Borrell *et al*. (2020); Rellstab *et al*. (2016)), does not assume that populations are currently at their adaptive optima but rather that overall GEAs across populations are generally adaptive. It quantifies the mean absolute deviation between a population’s observed genomic composition and the expected genomic composition derived from GEAs estimated for the target environment. Although both metrics are frequently interpreted as measures of maladaptation risk, their mathematical formulations, underlying assumptions, and biological meanings differ (Gain *et al*., 2023). The extent to which they provide comparable or consistent predictions remains poorly resolved.

GO estimates are also sensitive to the GEA methods employed (Archambeau *et al*., 2026; Fitzpatrick *et al*., 2021; Lind *et al*., 2024). This sensitivity may partly reflect differences in whether methods assume linear or non-linear genotype-environment relationships. Whether such relationships are linear or non-linear in nature remains an open question and method performance may depend on the underlying structure of the data. For instance, non-linear GEA methods may better capture sharp environmental gradients, whereas linear methods may be more appropriate for smooth gradients (Rellstab *et al*., 2021). Furthermore, the efficacy of these approaches is likely contingent upon the underlying genetic architecture of adaptation. Multivariate GEA approaches, for example, are generally expected to capture polygenic adaptation more effectively than univariate approaches (Forester *et al*., 2018). In addition, GEA methods used to identify climate-associated SNPs such as BayPass, LFMM, RDA, and partial RDA often detect different numbers and sets of candidate SNPs (Archambeau *et al*., 2026; Fitzpatrick *et al*., 2021; Lind & Lotterhos, 2025). These differences likely stem from variation in how each method accounts for population structure and how association thresholds are defined (e.g., p-value cutoffs, Bayes factors) (Forester *et al*., 2018; Fraņcois *et al*., 2016; Rellstab *et al*., 2015). Because GO is estimated using these identified SNPs, methodological differences in SNP detection may directly influence GO predictions. However, recent studies suggest that GO estimates derived from randomly selected SNP sets can perform similarly to those based on climate-associated SNPs (Fitzpatrick *et al*., 2021; Lachmuth *et al*., 2023; Lind *et al*., 2024), calling into question the necessity of SNP preselection.

Beyond that, GO implicitly assumes that current GEAs reflect adaptive relationships that remain stable across space and time, and that these associations can be extrapolated to unsampled environments across species ranges (i.e., populations with unknown genomic backgrounds) or future climatic conditions. However, if climate adaptation is polygenic, its genomic basis may differ among populations, as multiple genetic combinations can produce similar adaptive phenotypes (Capblancq *et al*., 2020a; Fitzpatrick *et al*., 2026; Rellstab *et al*., 2021). Furthermore, GEAs inferred under present conditions may not hold under novel environmental scenarios, particularly those involving unique combinations of climatic factors (Fitzpatrick *et al*., 2018, 2026; Lind & Lotterhos, 2025). Crucially, GO may not necessarily scale linearly with fitness decline; populations with similar or even lower GO values might experience greater fitness loss than those with higher estimates. This occurs because both allele frequency shifts and fitness responses are intimately dependent on the specific structure of environmental gradients and the underlying fitness landscape (Ahrens *et al*., 2026; Láruson *et al*., 2022). Consequently, empirical validation of GO estimates is necessary prior to their application in making conservation decisions.

Validation is commonly performed by testing for negative correlations between GO metrics and fitness components or proxies, with trait measurements from common garden experiments often used as proxies for fitness. While GO has generally been reported to predict fitness more effectively than climatic or geographic distance alone, results are not universally consistent, and its predictive ability may vary among species, fitness proxies, and life history stages (Archambeau *et al*., 2026; Capblancq & Forester, 2021; Fitzpatrick *et al*., 2021; Francisco *et al*., 2025; Gain *et al*., 2023; Ingvarsson & Bernhardsson, 2020; Lachmuth *et al*., 2023; Ĺaruson *et al*., 2022; Lind & Lotterhos, 2025; Rhońe *et al*., 2020; Verrico *et al*., 2026). A smaller number of studies have validated GO using long-term observational data from natural populations, finding that populations with higher GO tend to be declining more rapidly (Bay *et al*., 2018; Ruegg *et al*., 2018; Smith *et al*., 2021). In addition, GO is frequently evaluated using the same datasets employed to estimate GEAs, raising concerns about overfitting and inflated predictive performance (Capblancq *et al*., 2020b; Lotterhos, 2024). There is therefore a clear need for additional studies that assess GO across different species, methodologies, and realistic ecological contexts that encompass diverse environmental stressors, environmental gradients, and spatial contexts using independent, long-term observational data to clarify the relevance of GO in estimating the risk of maladaptation.

Here, we evaluate genomic offset (GO) as a predictor of maladaptation in North American butterflies of the genus *Lycaeides*. To date, most applications and empirical validations of GO have focused on long-lived taxa such as forest trees, which are characterized by long generation times and relatively slow evolutionary responses. In contrast, short-lived, highly fecund organisms such as insects remain underrepresented in GO validation studies, despite their well-documented capacity for rapid evolutionary responses (Garnas, 2018; Rudman *et al*., 2022; Simon & Peccoud, 2018). Such rapid evolutionary responses may buffer populations against abrupt environmental shifts, thereby altering genotype–environment associations over short timescales (Fitzpatrick *et al*., 2026; Rellstab *et al*., 2021). Consequently, GO may underestimate or fail to capture maladaptation risk in such populations. However, the extent to which GO remains a reliable predictor of maladaptation in species with life-history traits that facilitate rapid adaptation remains unclear. Addressing this gap is particularly urgent given the widespread, well-documented decline of insect populations driven by climate change and other anthropogenic pressures (Blüthgen *et al*., 2023; Edwards *et al*., 2025; Forister *et al*., 2021; Wagner *et al*., 2021). Butterflies provide a particularly powerful system for testing these ideas due to extensive genomic resources and long-term, spatially replicated demographic monitoring programs (Edwards *et al*., 2025; Henry *et al*., 2025; Taron & Ries, 2015).

The genus *Lycaeides*–which has been reclassified within the genus *Plebejus*, though we retain the historical nomenclature here to match prior literature and to precisely delineate our focal lineages (Talavera *et al*., 2013)–comprises a complex of several nominal species that are geographically widespread in North America, including *Lycaeides anna*, *L. idas*, *L. melissa*, and *L. samuelis* (Forister *et al*., 2011; Gompert *et al*., 2006; Nabokov, 1949). These species exhibit extensive variation across a range of ecological and life-history traits, including habitat use, phenology, voltinism, host plant use, mating behavior, and morphological characteristics such as wing patterning, egg morphology, and male genital morphology (Fordyce *et al*., 2002; Forister *et al*., 2006, 2009; Gompert *et al*., 2013b,a; Lucas *et al*., 2008, 2018; Nice & Shapiro, 1999). Reproductive isolation among these nominal species is incomplete, and past and ongoing hybridization have led to the formation of hybrid zones and admixed lineages characterized by combinations of parental, intermediate, and transgressive phenotypes (Chaturvedi *et al*., 2020; Gompert *et al*., 2006, 2010, 2012, 2013b; Nice *et al*., 2013). Recent work in *Lycaeides* used multivariate linear regression and genotyping-by-sequencing data to estimate GO and evaluate its relationship with demographic parameters inferred from genomic data rather than empirical changes in population abundance. Although no association was detected between GO and contemporary effective population size, GO was associated with genetic diversity and local rates of gene flow (Goodwin *et al*., 2026). However, it remains unknown whether GO predicts actual demographic responses, such as population declines or increases, and how sensitive these predictions are to the underlying methodological choices used to estimate GO.

We used pooled whole genome sequencing data from 29 *Lycaeides* populations, encompassing three nominal species (*L. anna*, *L. idas*, *L. melissa*) and admixed or hybrid lineages, together with long-term observational data from the North American Butterfly Association (NABA) in the western United States (Figure 1), to address three primary questions: (1) How robust are GO estimates to differences in SNP set composition and size across random SNP sets and climate-associated SNP sets identified by different GEA methods? (2) How consistent are GO estimates across different GO metrics and GEA methods? (3) How well does GO predict population trends, and how sensitive is its predictive performance to SNP set composition, GO metric, and GEA approach when evaluated using both training and independent datasets? The conceptual design of the study is illustrated in Figure 2.

**Figure 2:**
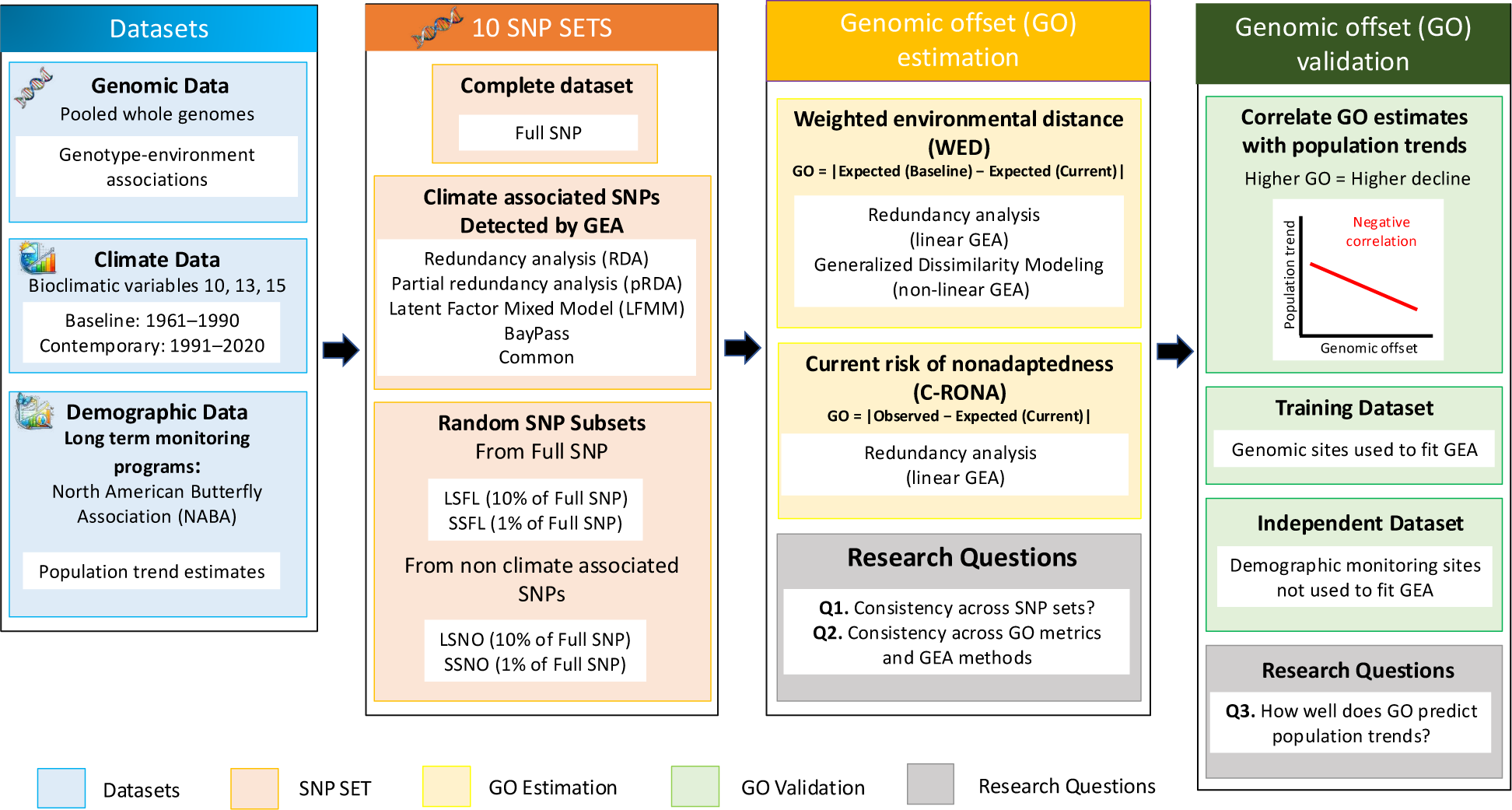
Workflow for addressing the three focal research questions. Genomic data from 29 *Lycaeides* populations were combined with baseline climate data (1961–1990) and contemporary climate data (1991–2020). Ten SNP sets were generated, the full SNP dataset, random subsets of the full dataset (large and small; LSFL and SSFL), climate-associated SNPs identified using four genotype–environment association (GEA) methods (pRDA, RDA, LFMM, BayPass), SNPs identified by all four methods (common SNPs), and random subsets of SNPs not identified as climate-associated by any method (large and small; LSNO and SSNO). These SNP sets were used to estimate two genomic offset (GO) metrics, weighted environmental distance (WED) and current risk of non-adaptedness (C-RONA), using redundancy analysis (RDA) and generalized dissimilarity modeling (GDM). The first two research questions examined consistency of GO estimates across SNP sets, GEA methods, and GO metrics. The third question validated the predictive performance of GO using multidecadal demographic data from NABA monitoring sites. Arrows indicate data flow into each analysis.

## Materials and Methods

### Genomic dataset

#### DNA extraction and sequencing

DNA was extracted from 1358 butterflies sampled from 29 *Lycaeides* populations in the western United States using the Qiagen DNeasy 96 Blood and Tissue Kit (Table S1, Figure 1c). The DNA concentration for each individual was quantified using a fluorescence microplate reader (Turner Biosystems Modulus Microplate). Equal amounts of DNA from individuals within each population were pooled for whole genome sequencing.

Libraries were prepared and sequenced by BGI using DNB-based sequencing technology on the DNBseq platform (paired-end 100 bp), targeting approximately 100× coverage per population. Raw reads were quality-filtered using SOAPnuke (Chen *et al*., 2018) and aligned to the *L. melissa* reference genome using bwa-mem2 (v2.0pre2) (Li & Durbin, 2009; Vasimuddin *et al*., 2019). PCR duplicates were removed with SAMtools (v1.16) (Ebbert *et al*., 2016; Li *et al*., 2009). SNPs were called using the bcftools consensus caller (-c; v1.16) (Li, 2011) and subsequently filtered with GATK (v4.1.4.1) (McKenna *et al*., 2010). Full details of library preparation, read filtering thresholds, alignment parameters, variant calling, and SNP filtering criteria are provided in the Supplementary Methods.

For each SNP, read counts supporting reference and alternate alleles were extracted. These counts were used in R (v4.2.2) to estimate alternate allele frequencies as the ratio of alternate read counts to total depth per locus. SNPs missing allele frequency estimates in any population and SNPs exhibiting no allele frequency variation among populations were excluded. The final genomic dataset comprised 16,285,847 SNPs, which were used in all downstream analyses.

### Climatic dataset

Climate data were extracted from the Climate Downscaling Tool (ClimateDT v.2.0) using the bilinear interpolation method (Marchi *et al*., 2024) for both the 29 *Lycaeides* populations sampled for genomic analyses and the independent long-term North American Butterfly Association (NABA) monitoring locations where *Lycaeides* butterflies have been surveyed annually (Figure 1c).

We extracted data for bioclimatic variables (Bio1–Bio19) for baseline and contemporary periods. Climate data from 1961–1990 were used as the baseline climate, consistent with prior ecological and genomic offset studies, as this period precedes substantial global warming (IPCC, 2014; Lachmuth *et al*., 2023, 2024; Lind *et al*., 2024; Wang *et al*., 2016). Climate data from 1991–2020 were used as contemporary climate for validation analyses. For both periods, climatic variables were averaged across years for each location.

To identify the subset of climatic variables that best explained genetic variation among populations sampled for genomic analysis, we implemented a RDA-based stepwise variable selection procedure following Capblancq *et al*. (2018) and Capblancq & Forester (2021). Variable selection was conducted using the ordiR2step function of the vegan package (v2.6-10) in R. Selection followed the double stopping criterion of Blanchet *et al*. (2008), whereby variables were added sequentially based on 1,000 permutations and the procedure stopped when additional variables were no longer significant (*α* ≥ 0.01) or when the adjusted *R*^2^ of the reduced model equaled or exceeded that of the full model. This procedure identified a best model with bioclimatic variables 10 (Mean Temperature of Warmest Quarter), 13 (Precipitation of Wettest Month), and 15 (Precipitation Seasonality), which were used in all downstream analyses.

### Ten SNP sets

To evaluate how SNP set composition and size influence GO estimates, we generated ten SNP sets representing three broad categories: (i) the complete SNP dataset, (ii) five climate-associated SNP sets, including four sets identified by individual GEA methods and one set comprising SNPs detected by all four methods, and (iii) four randomly sampled SNP sets (Table 1).

**Table 1:**
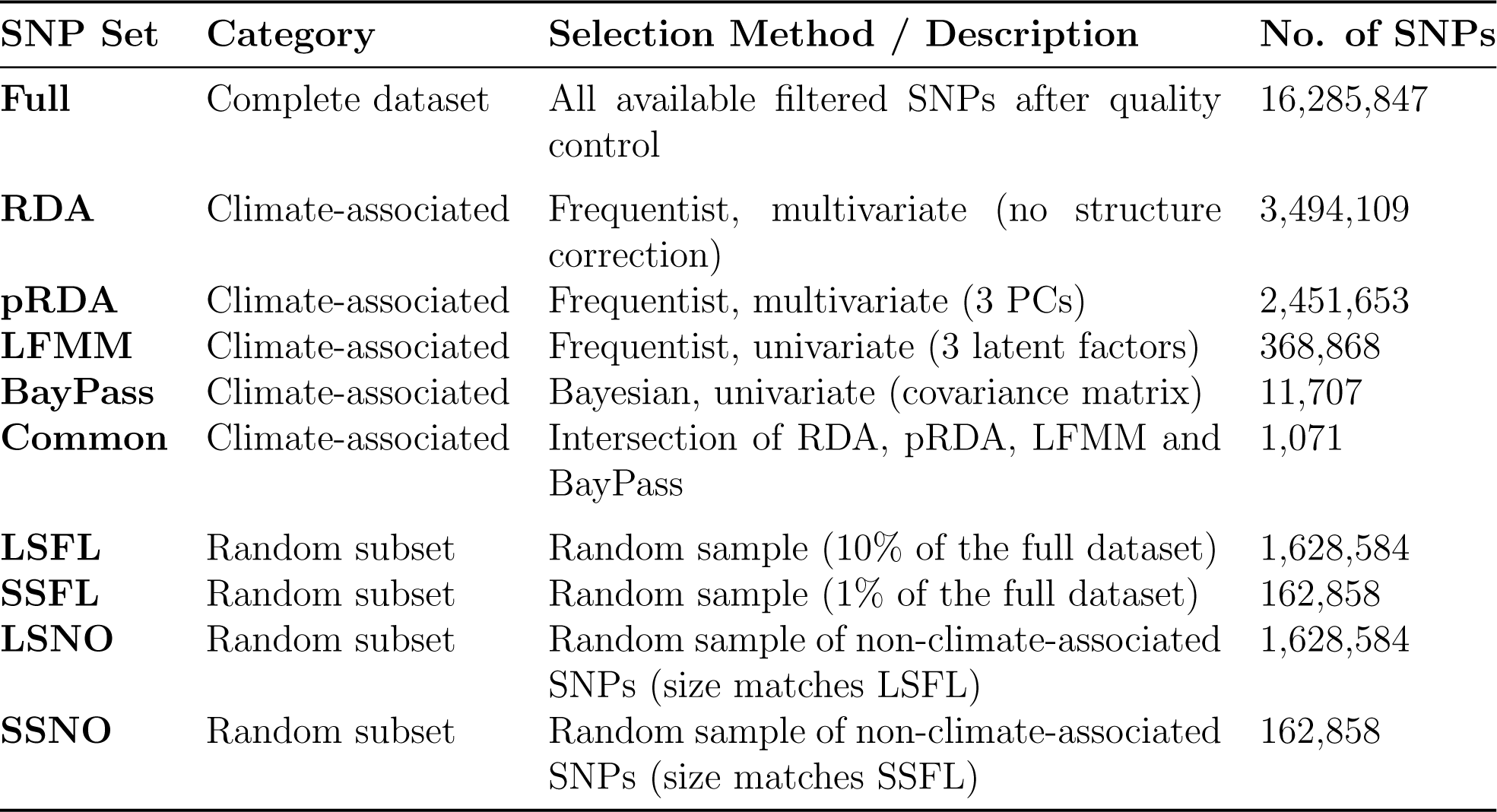
Summary of the ten SNP sets used to evaluate GO estimates. LSFL and SSFL represent 10% and 1% of the full dataset, respectively, while LSNO and SSNO represent equivalently sized samples drawn exclusively from non-climate-associated SNPs.

| **SNP Set** | **Category** | **Selection Method / Description** | **No. of SNPs** |
| --- | --- | --- | --- |
| **Full** | Complete dataset | All available filtered SNPs after quality control | 16,285,847 |
| **RDA** | Climate-associated | Frequentist, multivariate (no structure correction) | 3,494,109 |
| **pRDA** | Climate-associated | Frequentist, multivariate (3 PCs) | 2,451,653 |
| **LFMM** | Climate-associated | Frequentist, univariate (3 latent factors) | 368,868 |
| **BayPass** | Climate-associated | Bayesian, univariate (covariance matrix) | 11,707 |
| **Common** | Climate-associated | Intersection of RDA, pRDA, LFMM and BayPass | 1,071 |
| **LSFL** | Random subset | Random sample (10% of the full dataset) | 1,628,584 |
| **SSFL** | Random subset | Random sample (1% of the full dataset) | 162,858 |
| **LSNO** | Random subset | Random sample of non-climate-associated SNPs (size matches LSFL) | 1,628,584 |
| **SSNO** | Random subset | Random sample of non-climate-associated SNPs (size matches SSFL) | 162,858 |

#### Climate-associated SNPs

Climate-associated SNPs were identified with four widely used GEA methods: LFMM, BayPass, RDA, and pRDA (hereafter referred to as LFMM SNPs, BayPass SNPs, RDA SNPs, and pRDA SNPs). These methods differ in statistical framework (Bayesian vs. frequentist), dimensionality (univariate vs. multivariate), and whether they account for population structure (Table S2). Baseline climate data from 1961–1990, representing the conditions to which populations are assumed to be locally adapted, were used as predictor variables in all analyses.

Population structure was accounted for using the first three principal components derived from centered and scaled SNP allele frequencies in pRDA and as three latent factors in LFMM (Table S1a). This choice ensured a consistent representation of major axes of genetic structure across both methods. BayPass accounted for population structure through an estimated covariance matrix of allele frequencies among populations, while RDA did not explicitly correct for population structure (Archambeau *et al*., 2026; Capblancq & Forester, 2021; Gautier, 2015).

We further evaluated whether additional correction for spatial autocorrelation was required using distance-based Moran’s Eigenvector Maps. This indicated that spatial structure was largely redundant with population structure and did not provide substantial independent explanatory power (Table S3). Accordingly, we did not correct spatial autocorrelation in our core analyses.

Finally, we identified SNPs detected as climate-associated by all four methods, hereafter called “Common SNPs”. SNPs identified by each individual method, as well as the Common SNPs, were retained for downstream analyses. Detailed descriptions of each GEA method, including model specifications, parameters, association thresholds, and evaluation of spatial autocorrelation, are provided in the Supplementary Methods.

#### Random Selection of SNPs

We randomly sampled 1% (162,858 SNPs) and 10% (1,628,584 SNPs) of the full SNP dataset (16,285,847 SNPs). The same subset sizes were then applied to the SNPs not identified as climate-associated by any GEA method (hereafter non-climate-associated SNPs), ensuring consistency in SNP number and allowing for direct comparison across subsets of similar size. This yielded four random subsets: a large random sample of the full dataset (LSFL; n = 1,628,584 SNPs), a small random sample of the full dataset (SSFL; n = 162,858 SNPs), a large random subset of non-climate-associated SNPs (LSNO; n = 1,628,584 SNPs), and a small random subset of non-climate-associated SNPs (SSNO; n = 162,858 SNPs).

### Genomic offset estimation

We estimated the current risk of nonadaptedness (C-RONA) and weighted environmental distance (WED) using two multivariate GEA approaches, RDA and generalized dissimilarity modeling (GDM). RDA assumes linear relationships, whereas GDM models non-linear associations. These approaches were selected because multivariate GEA methods are better suited for detecting polygenic adaptation, which is most likely the case for adaptation to climate (Candido-Ribeiro *et al*., 2026; R^ego *et al*., 2025). Gradient forest was not used due to computational constraints associated with the size of the SNP datasets. For each of the ten SNP sets (full SNP dataset, pRDA SNPs, RDA SNPs, LFMM SNPs, BayPass SNPs, Common SNPs, LSFL, SSFL, LSNO, SSNO) we estimated C-RONA and WED using RDA and WED using only GDM. This is because C-RONA requires residuals for each population, whereas GDM produces residuals for pairwise population dissimilarities, not per-population values, making it unsuitable for C-RONA estimation. GDM and RDA were implemented in R using the gdm and vegan packages, respectively.

#### Weighted environmental distance (WED) estimation

WED was estimated using RDA following Capblancq & Forester (2021) and using GDM following Fitzpatrick & Keller (2015). For RDA, models were fitted using baseline climate as predictors and SNP allele frequencies as response variables. WED was calculated as the Euclidean distance between expected allele frequencies under baseline and contemporary climate. GDM models non-linear associations between pairwise genetic distance and geographic and environmental distance among populations. GDM was fitted using pairwise F_ST_ matrices (29 × 29) derived from each SNP set as response variables. Predictor variables consisted of pairwise geographic distance and multivariate baseline climatic distance. After fitting the model, climatic distances between baseline and contemporary climate were used to predict WED, representing expected genetic distance between population allele frequencies under baseline climate and population allele frequencies under contemporary climate.

#### Current risk of nonadaptedness (C-RONA) estimation

Borrell *et al*. (2020) estimated C-RONA using linear regressions, with climate as predictor variables and SNP data as response variables. C-RONA was calculated as the mean absolute difference between observed allele frequencies and those expected under the fitted GEA model. Here, we estimate C-RONA using RDA. For each SNP set, RDA was performed with contemporary climate as predictor variables and SNP allele frequencies as response variables. C-RONA was then calculated as the mean absolute residual between observed allele frequencies and those predicted by the fitted GEA model for each populations sampled for genomic analysis.

#### Sensitivity of GO estimates to SNP set composition and size

To address our first question regarding whether GO estimates are similar across random SNP sets and climate-associated SNPs identified by different GEA methods, despite differences in composition and size, we performed pairwise Spearman correlations of GO estimates across populations sampled for genomic analysis using each SNP set. We conducted this analysis separately for RDA-based C-RONA, RDA-based WED, and GDM-based WED.

#### Consistency of GO estimates across GO metrics and GEA methods

To address our second question examining whether GO estimates vary by GO metric and GEA method, we performed pairwise Spearman correlations between GO estimates across populations sampled for genomic analysis. We compared RDA-based C-RONA and RDA-based WED (comparison of GO metrics), RDA-based WED and GDM-based WED (comparison of GEA methods for estimating GO), and RDA-based C-RONA and GDM-based WED (comparison of both GO metrics and GEA methods). These correlations were performed for each SNP set.

### Predictive performance of GO across SNP sets, GO metrics, and GEA approaches

To evaluate how well GO predicts population trends and how sensitive its predictive performance is to SNP set composition, GO metric, and GEA approach, we validated GO using both training and independent datasets. We evaluated GO at sites where genomic data were used to estimate GEA relationships (training dataset) and at NABA monitoring sites where genomic data were not used to estimate GEA relationships (independent dataset) (Figure 1c). We estimated population trends of *Lycaeides* butterfly populations at monitored sites using the NABA dataset. Population trends at each monitoring site were estimated using hierarchical Bayesian models that account for site-specific effects. In this dataset, the temporal effect of year was used to estimate population trends, with negative values indicating declining populations and positive values indicating increasing populations. Trends at populations sampled for genomic analysis were subsequently derived through spatial interpolation from nearby monitored sites. Full details of the dataset, modeling framework, priors, posterior sampling, and interpolation procedures are provided in the Supplementary Methods.

#### Training dataset validation

For the training dataset validation, we quantified the association between GO estimates and population trends across populations sampled for genomic analysis using Spearman correlations for each SNP set. To benchmark GO performance, we also calculated the Euclidean distance between baseline and contemporary climate conditions for each populations sampled for genomic analysis and assessed its relationship with population trends using Spearman correlations.

#### Independent dataset validation

We conducted an analogous evaluation using the independent NABA dataset. In this case, GEA relationships estimated from the populations sampled for genomic analysis using both RDA and GDM were applied to predict WED for each monitored site based on their baseline and contemporary climate conditions. C-RONA was not estimated for monitored sites, as it requires observed allele frequencies which are unavailable for this dataset. For each SNP set, we calculated Spearman correlations between GO estimates and population trends across monitored sites. As with the training sites, we also assessed correlations between climate distance and population trends to facilitate direct comparison with GO estimates. In both analyses, negative correlations were expected, reflecting steeper population declines in populations with higher genomic offsets.

## Results

### Identifying climate-associated SNPs

Climate-associated SNPs were designated using LFMM, pRDA, RDA, and BayPass. We observed substantial variability in both the number and identity of climate-associated SNPs detected by each method. Specifically, LFMM identified 368,868 SNPs (*q <* 0.05), pRDA identified 2,451,653 SNPs (*q <* 0.05), RDA identified 3,494,109 SNPs (*q <* 0.05), and BayPass identified 11,707 climate-associated SNPs (*Bayesfactor >* 20). Multivariate GEA approaches (RDA and pRDA) detected a substantially larger number of SNPs compared to univariate approaches. Only 1,071 SNPs were identified as climate-associated by all four GEA methods (Figure 3a).

**Figure 3:**
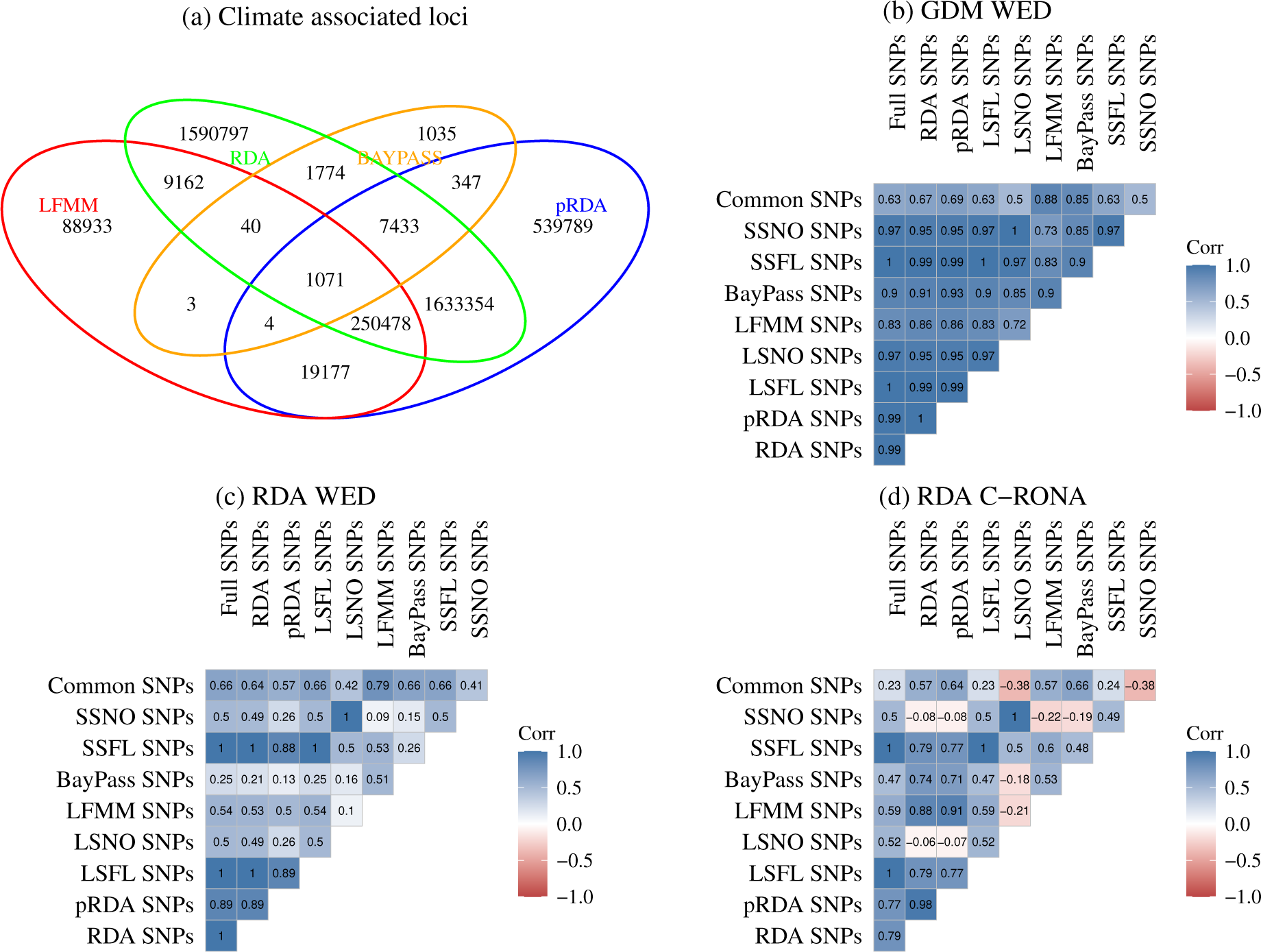
Distribution of climate-associated loci and consistency of genomic offset (GO) estimates across SNP sets. (a) Venn diagram showing the number of climate-associated SNPs identified by each genotype–environment association (GEA) method and the overlap among methods. (b–d) Pairwise Spearman correlations of GO estimates across ten SNP sets for populations at each genomic sampling site. GO was estimated as (b) weighted environmental distance (WED) using generalized dissimilarity modeling (GDM), (c) WED using redundancy analysis (RDA), and (d) current risk of non-adaptedness (C-RONA) using RDA. The ten SNP sets include the full SNP dataset (Full SNPs), large and small random samples of the full dataset (LSFL and SSFL), climate-associated SNPs identified by four GEA methods (pRDA, RDA, LFMM, BayPass), SNPs identified by all methods (Common SNPs), and large and small random subsets of non climate-associated SNPs (LSNO and SSNO). Positive correlation coefficients indicate similarity in GO estimates between compared SNP sets.

### Sensitivity of GO estimates to SNP set composition and size

We examined the consistency of GO estimates across SNP sets varying in composition and size by calculating pairwise Spearman correlations (*ρ*) of GO estimates (Figure 1c, S2-S4) for each populations sampled for genomic analysis across all SNP sets. This analysis was conducted separately for RDA-based C-RONA, RDA-based WED, and GDM-based WED estimates. For WED estimates, we found moderate positive correlations (*ρ* ≥ 0.40) between GO estimates across SNP sets differing in both composition and size (Figure 3b,c). However, exceptions were observed. Correlations among RDA-based WED estimates were weak when using pRDA, LFMM or BayPass SNPs. In contrast, C-RONA estimates exhibited a wide range of correlations (-0.38 ≤ *ρ* ≤ 1.00). Notably, negative correlations (-0.38 ≤ *ρ* ≤ −0.06) were observed between GO estimates derived from random non-outlier SNPs and those from SNPs identified as climate-associated by one or more GEA methods (Figure 3d). Overall, while GO estimates are generally consistent across SNP sets, the degree of consistency depends on the GO metric and the SNP set used.

### Consistency of GO estimates across GO metrics and GEA approaches

We assessed the sensitivity of GO estimates to the choice of GO metric and GEA method by calculating pairwise Spearman correlations between GO estimates for each populations sampled for genomic analysis. We compared: (1) RDA-based C-RONA versus RDA-based WED to evaluate consistency between GO metrics, (2) RDA-based WED versus GDM-based WED to evaluate consistency between GEA methods, and (3) RDA-based C-RONA versus GDM-based WED to evaluate consistency when both GO metric and GEA method differ. Correlations were calculated separately for each SNP set (Figure 4a). When comparing GEA methods (RDA-based WED and GDM-based WED), correlations depended on the SNP set. Significant positive correlations were observed for Full (*ρ* = 0.38, 95% CI: 0.01-0.68), pRDA (*ρ* = 0.51, 95% CI: 0.21-0.73), LSFL (*ρ* = 0.38, 95% CI: 0.01-0.68), and SSFL SNPs (*ρ* = 0.37, 95% CI: 0.02-0.67). These results indicate that the similarity between GO estimates derived from different GEA methods is sensitive to SNP set composition, but not necessarily related to SNP set size. In contrast, comparisons between GO metrics (RDA-based WED and RDA-based C-RONA) revealed non-significant correlations across all SNP sets. This lack of consistency suggests that C-RONA and WED yield substantially different estimates of maladaptation risk for the same populations, and thus measure different things. Finally, when comparing both GO metrics and GEA methods simultaneously (RDA-based C-RONA and GDM-based WED), significant positive correlations were observed for SNP sets not specifically detected as climate-associated by any GEA method: Full SNPs (*ρ* = 0.39, 95% CI: 0.01–0.67), LSFL (*ρ* = 0.39, 95% CI: 0.003–0.67), LSNO (*ρ* = 0.65, 95% CI: 0.35–0.82), SSFL (*ρ* = 0.38, 95% CI: 0.01–0.67), and SSNO (*ρ* = 0.66, 95% CI: 0.34–0.83). Again, correlation strength varied by SNP set (Figure 4a).

**Figure 4:**
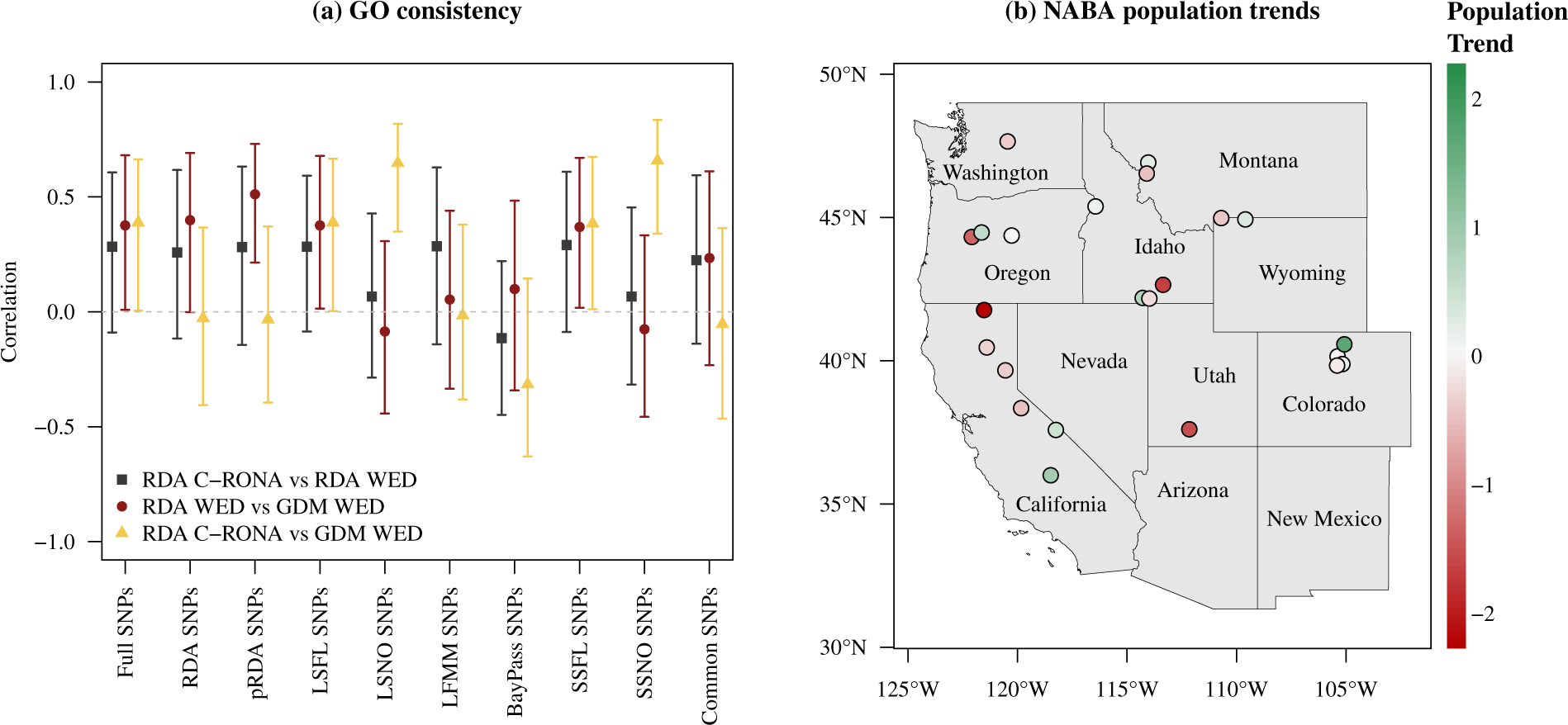
Consistency of genomic offset (GO) estimates across GO metrics and genotype–environment association (GEA) methods and spatial distribution of population trends. (a) Consistency of genomic offset (GO) estimates across GO metrics and GEA methods for each SNP set. Pairwise Spearman correlations compare: (1) RDA-based current risk of non-adaptedness (C-RONA) versus RDA-based weighted environmental distance (WED) to evaluate consistency between GO metrics; (2) RDA-based WED versus GDM-based WED to evaluate consistency between GEA methods; and (3) RDA-based C-RONA versus GDM-based WED to evaluate consistency when both GO metric and GEA method differ. SNP sets are ordered from left to right by decreasing number of SNPs (Full SNPs contains the most SNPs and Common SNPs the fewest). Error bars denote 95% confidence intervals. Significant positive correlations indicate greater similarity in GO estimates. (b) Spatial distribution of population trends in *Lycaeides* populations across the western United States, estimated from the North American Butterfly Association (NABA) dataset. Each point represents a monitoring site, with color indicating the estimated population trend. Red denotes declining abundance (negative trend), white indicates stable populations, and green denotes increasing abundance (positive trend).

### Predictive performance of GO across SNP sets, GO metrics, and GEA approaches

We estimated population trends for *Lycaeides* populations at monitored sites using the NABA dataset. Our analyses revealed substantial spatial heterogeneity in population trends across the western United States, although declines were more widespread than increases (Figure 4b). We then evaluated how well GO predicts population trends and how sensitive its predictive performance is to SNP set composition, GO metric, and GEA approach by quantifying associations between GO estimates (Figure S2-S6) and population trends (Figure 4b) using Spearman rank correlations. We expected negative correlations, reflecting steeper population declines in populations experiencing higher genomic offset. GO was validated using both the populations used to estimate GEAs (hereafter, “training dataset”) and the demographic monitoring sites, which were not part of the GEA analysis (hereafter “independent datasets”) (Figure 5).

**Figure 5:**
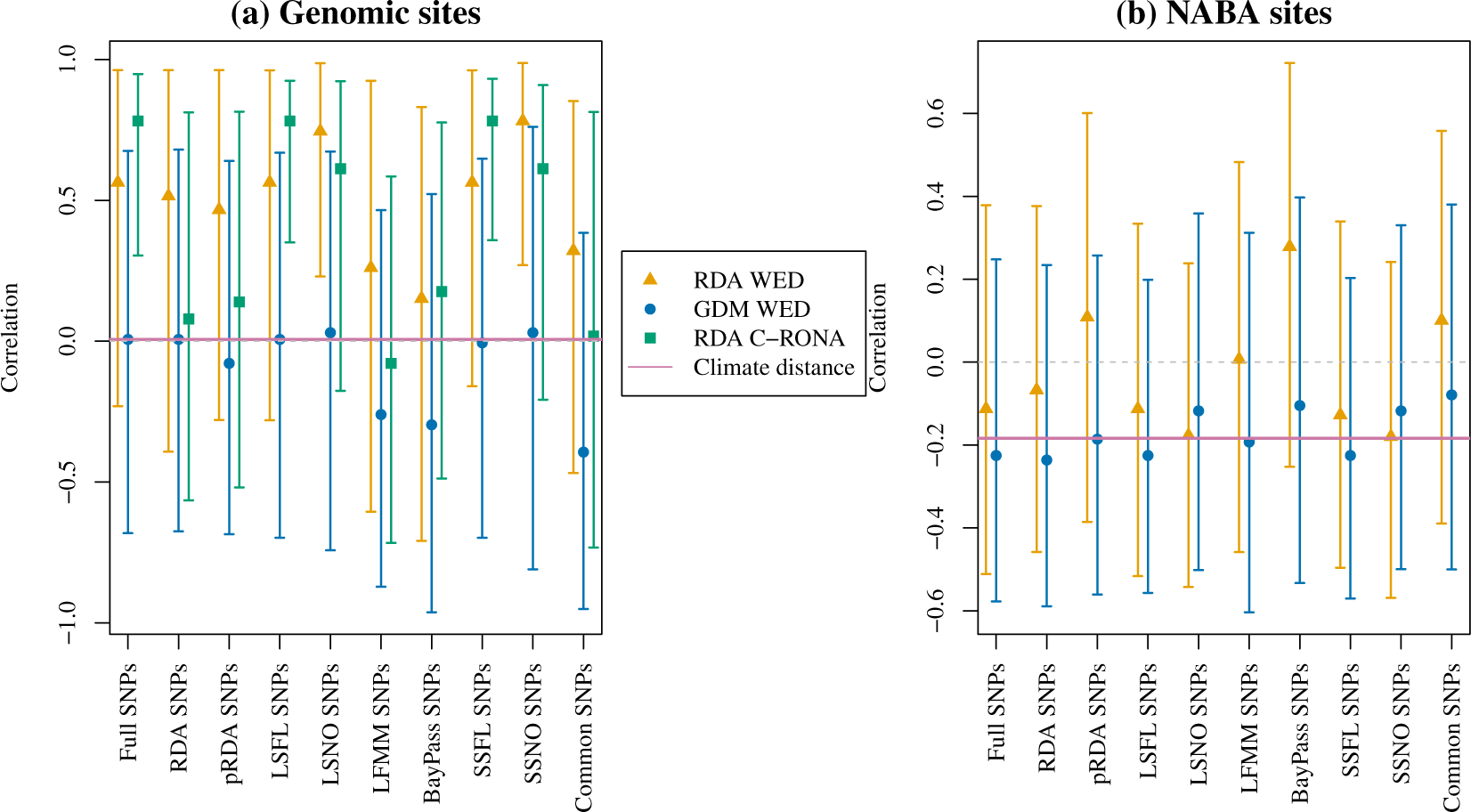
Validation of the predictive performance of genomic offset (GO) across SNP sets, GO metrics, and genotype–environment association (GEA) methods. Spearman rank correlations between GO estimates and observed population trends were calculated to assess predictive performance. (a) Training dataset validation: GO estimates were evaluated at populations sampled for genomic analysis used to fit GEA relationships. Population trends at these sites were estimated via spatial interpolation from nearby North American Butterfly Association (NABA) monitoring sites. Correlations are shown for RDA-based weighted environmental distance (WED), GDM-based WED, and RDA-based current risk of non-adaptedness (C-RONA) across all SNP sets. (b) Independent dataset validation: GEA relationships estimated from populations sampled for genomic analysis were extrapolated to predict GO at NABA monitoring sites not used to fit GEAs. Correlations between GO estimates (RDA-based WED and GDM-based WED) and observed population trends are shown for all SNP sets. SNP sets are ordered from left to right by decreasing number of SNPs. Error bars denote 95% confidence intervals. Significant negative correlations would indicate that GO is predictive of population trends (higher GO associated with steeper declines).

For the training dataset (Figure 5a), GO estimates generally showed weak and non-significant correlations with population trends, regardless of SNP set composition or size, GO metric, or GEA approach. Contrary to expectations, significant positive correlations were observed between population trends and RDA-based C-RONA estimates for Full (*ρ* = 0.78, 95% CI: 0.30-0.95), LSFL (*ρ* = 0.78, 95% CI: 0.35-0.92), and SSFL (*ρ* = 0.78, 95% CI: 0.36-0.93) as well as correlations between population trends and RDA-based WED estimates for both LSNO (*ρ* = 0.75, 95% CI: 0.23-0.99), and SSNO (*ρ* = 0.78, 95% CI: 0.27-0.99), indicating that populations with higher genomic offset exhibited more positive population trends. Similarly, climate distance (*ρ* = 0.01, 95% CI: -0.74-0.70) showed weak, non-significant correlations with population trends. These results suggest that GO is poorly predictive of population trends and may perform particularly poorly when estimated using SNPs not detected as climate-associated by GEA methods.

For the independent dataset (Figure 5b), we observed similar patterns, with GO estimates showing non-significant correlations with population trends regardless of SNP set composition, SNP set size, or GEA approach. Likewise, climate distance was not significantly correlated with population trends (*ρ* = −0.18, 95% CI: -0.53-0.27). Overall, our results suggest that GO is a poor predictor of recent demographic trends in *Lycaeides* butterflies.

## Discussion

Genomic offset (GO) is increasingly used to estimate the risk of maladaptation under climate change, with populations exhibiting higher GO assumed to face greater risk. Despite its growing application in conservation planning, critical questions remain about how reliably GO estimates reflect biological reality, and how this depends on the choice of SNP set, GO metric, and GEA method used to estimate it. Empirical validation against observed population responses is also rarely performed. Here, using *Lycaeides* butterflies, a well studied insect with short generation times and high fecundity that may confer capacity for rapid adaptation (Garnas, 2018; Simon & Peccoud, 2018), we provide a comprehensive evaluation of GO consistency and predictive performance across multiple analytical frameworks.

Consistent with previous studies by Archambeau *et al*. (2026) and Fitzpatrick *et al*. (2021), we found that the identity and number of climate-associated SNPs varied substantially among GEA methods. Multivariate approaches (RDA and pRDA) detected a larger number of SNPs compared to univariate methods (LFMM, BayPass), but only a small set of SNPs was consistently identified across all methods. Despite this variability, GO estimates derived from SNPs identified by GEA methods were largely consistent with those derived from randomly selected SNPs, suggesting that identifying climate-associated SNPs prior to estimating GO may not substantially alter inference. Similar findings have been reported by Fitzpatrick *et al*. (2021), Ĺaruson *et al*. (2022), Lind *et al*. (2024), and Lind & Lotterhos (2025), who found that climate-associated SNPs detected by GEA methods do not outperform randomly selected SNPs in GO estimation. This pattern likely reflects the pervasive confounding of population structure and climatic gradients. When population structure is strongly correlated with environmental variation, virtually all loci, not only those under direct climatic selection will carry signal correlated with climate. Under these conditions, it becomes difficult to isolate a signal of climate adaptation from the broader genomic background, a challenge that is particularly relevant for studies seeking to identify genomic regions important for climate adaptation (Booker, 2024; Lotterhos, 2023).

The consistency of GO estimates across GEA methods depended on the SNP set analyzed. While some SNP sets yielded strong positive correlations between GO estimates derived from linear (RDA) and non-linear (GDM) methods, others showed weaker or inconsistent patterns. This finding aligns with simulation based work by Gain *et al*. (2023) and Rellstab *et al*. (2021), who showed that predictions from different GEA approaches can converge under certain conditions, and with empirical comparisons by Archambeau *et al*. (2026) and Lind *et al*. (2024), who reported variable consistency across GEA methods and species. Together, these results suggest that the choice of GEA method introduces context dependent uncertainty into GO estimation.

More strikingly, we found a consistent lack of agreement between the two GO metrics examined (WED and C-RONA) across all SNP sets. This finding is consequential because WED and C-RONA are often treated as interchangeable in the literature, yet they differ fundamentally in their mathematical formulations, biological assumptions, and interpretations (Gain *et al*., 2023). WED measures the environmental distance a population must traverse to reach a new adaptive optimum, weighted by the strength of genotype–environment associations, and assumes local adaptation without relying directly on observed allele frequencies of individual populations (Capblancq *et al*., 2020b; Capblancq & Forester, 2021; Fitzpatrick & Keller, 2015). C-RONA, by contrast, does not assume local adaptation of each population independently but treats observed allele frequencies as reflective of the overall GEA landscape. It estimates how far a population’s allele frequencies deviate from those expected under prevailing climate (Rellstab *et al*., 2016, 2021). Because these metrics capture different dimensions of maladaptation risk, their non-interchangeability is not merely a statistical concern but a biological one. Conclusions about which populations are most vulnerable may differ depending on which metric is used. Our results underscore the need for clearer theoretical frameworks that specify which metric, and which underlying assumptions, are most appropriate given the biology of the focal species and the conservation question being addressed.

GO was poorly predictive of population trends of *Lycaeides* butterflies, regardless of SNP set, GEA method, or GO metric. This held true whether validated at populations sampled for genomic analysis used to estimate GEA relationships (training dataset) or unsampled NABA sites where GEA relationships were extrapolated spatially (independent dataset). In some cases, correlations were contrary to the expected negative relationship, with higher GO associated with more positive population trends. Although spatial interpolation of demographic responses across heterogeneous landscapes can introduce uncertainty, as nearby populations may exhibit divergent trajectories due to fine-scale environmental variation (Nice *et al*., 2019; Reis *et al*., 2026), this is unlikely to account for our results, given that GO also showed poor predictive performance at monitored NABA sites. This lack of demographic predictability aligns with and extends the findings of Goodwin *et al*. (2026), who detected no relationship between GO and contemporary effective population size in this system. Our results contrast with findings from studies validating GO in forest trees and other long-lived species, where negative correlations have been reported between GO and fitness-related traits measured in common garden experiments (Archambeau *et al*., 2026; Capblancq & Forester, 2021; Fitzpatrick *et al*., 2021; Francisco *et al*., 2025; Gain *et al*., 2023; Rhońe *et al*., 2020), as well as with studies linking GO to expected population trends in birds (Bay *et al*., 2018; Ruegg *et al*., 2018). Climate distance, used as a simpler benchmark predictor, performed no better than GO, suggesting the weak signal is not specific to genomic approaches but may reflect broader challenges in linking any measure of climatic distance to demographic outcomes in this system.

Several non-mutually exclusive explanations may account for these findings. First, neutral demographic history may confound GEAs, as population structure often generates spurious signals that do not reflect adaptive variation (Forester *et al*., 2018; Láruson *et al*., 2022). This issue is exacerbated in small or genetically depauperate populations, where stochastic processes such as genetic drift can dominate allele frequency dynamics and constrain adaptive potential, potentially leading to low GO estimates even in populations at high risk of extirpation (Ahrens *et al*., 2026; Láruson *et al*., 2022; Van Deurs *et al*., 2025). Second, the predictability of GO may be influenced by the genetic architecture of adaptation. If climate adaptation is polygenic, with different genomic regions underlying local adaptation across populations, or if multiple allelic combinations produce similar adaptive phenotypes across populations (i.e., genetic redundancy), GEA analyses may fail to capture locally relevant adaptive loci. Under these conditions, SNPs identified as important for local adaptation in one set of populations may have limited relevance for predicting maladaptation in others, weakening the relationship between GO and observed demographic responses (Candido-Ribeiro *et al*., 2026; Láruson *et al*., 2020; Lotterhos, 2023; Yeaman, 2015).

Third, GO relies on the assumption that genetic distance scales linearly with fitness decline. Fitness responses depend on the complex structure of environmental gradients and underlying fitness landscapes (Exposito-Alonso *et al*., 2019; Mathieson & McVean, 2013; Simon & Coop, 2024; Trujillo *et al*., 2022; Van Deurs *et al*., 2025). For example, intermediate clinal populations may exhibit high GO due to steep allele frequency shifts, whereas populations at environmental extremes may show low GO yet still experience severe fitness declines as they exceed hard physiological limits (Angert *et al*., 2020; Bush *et al*., 2016; Hargreaves *et al*., 2014; Lee-Yaw *et al*., 2016; Moŕan-Ordóñez *et al*., 2018; Nadeau & Urban, 2019).

Fourth, life history may play an important role. Unlike the long-lived tree species in which GO has most often been validated, *Lycaeides* butterflies have short generation times and high fecundity, which may facilitate rapid adaptation. We hypothesize that these life history characteristics may attenuate the demographic consequences of climate maladaptation: if populations can track environmental change through rapid evolutionary responses, GO estimated from historical GEA relationships may not accurately reflect current maladaptation risk and may therefore fail to predict observed population trends (Garnas, 2018; Rudman *et al*., 2022; Simon & Peccoud, 2018). Broader comparative studies across species with contrasting life histories and adaptive capacities will be necessary to evaluate whether GO predictability is systematically related to generation time or evolutionary potential. Finally, inherent noise in broad-scale NABA monitoring data may obscure true demographic trajectories, masking any underlying relationship with GO. These findings have clear implications for conservation applications of GO. First, the choice of SNP set, GEA method, and GO metric can substantially influence estimates, and relying on a single approach may be misleading. Second, GO predictions should be validated against empirical population or fitness data whenever possible before using them for management decisions. Third, careful attention should be paid to the assumptions underlying each GO metric and GEA method, as well as species-specific factors such as demographic history, life history, and genetic architecture. More broadly, there is a pressing need for a deeper theoretical understanding of GO and its relationship to population viability under climate change. Conservation practitioners typically require information about whether populations will persist–a question of mean absolute fitness–yet it remains unclear whether GO actually reflects this. Bridging this gap will require both improved theory linking genomic maladaptation to demographic outcomes and empirical studies that explicitly test the assumptions of GO across species with diverse life histories, genetic architectures, and demographic contexts. To date, validation studies have been disproportionately focused on long-lived trees and a handful of other taxa; broader comparative validation across species with differing capacities for rapid adaptation will be essential for establishing the conditions under which GO can reliably guide conservation decisions. In conclusion, while GO may provide a useful conceptual framework for assessing potential maladaptation in some systems, our results underscore the need for careful application, rigorous validation, and a nuanced understanding of its underlying assumptions and scope of applicability.

## Acknowledgments

We gratefully acknowledge support from the U.S. National Science Foundation (DEB-2114973 to M.L.F., DEB-1844941 and DEB-2114974 to Z.G.), the Utah State University Ecology Center Research Award, the Society for the Study of Evolution R. C. Lewontin Early Award, and the American Philosophical Society Lewis and Clark fund for exploration and field research (to G.A.R.). We also acknowledge the computational resources provided by the Center for High Performance Computing at the University of Utah. We thank the members of the Gompert Lab at Utah State University for their valuable feedback on earlier drafts of this manuscript.

## Author Contributions

Gbolahan A Reis: Conceptualization (equal); Data Curation (equal); Formal Analysis (lead); Funding Acquisition (equal); Methodology (equal); Visualization (lead); Writing – Original Draft Preparation (lead); Writing – Review & Editing (equal). Lauren K Lucas: Data Curation (equal); Investigation (equal); Writing – Review & Editing (equal). Matthew L Forister: Data Curation (equal); Investigation (equal); Funding Acquisition (equal); Writing – Review & Editing (equal). Arthur M Shapiro: Data Curation (equal); Investigation (equal); Writing – Review & Editing (equal). James A Fordyce: Data Curation (equal); Investigation (equal); Writing – Review & Editing (equal). Chris C Nice: Data Curation (equal); Investigation (equal); Writing – Review & Editing (equal). Zachariah Gompert: Conceptualization (equal); Data Curation (equal); Funding Acquisition (equal); Methodology (equal); Supervision (lead); Writing – Review & Editing (equal).

## Data and code availability

Pooled whole-genome sequence data for *Lycaeides* have been deposited in the NCBI Sequence Read Archive (PRJNA1375794). All analytical code used in this study is available for review on GitHub (https://github.com/Reisanthony/Genomic offset manuscript and https://github.com/zgompert/LycAdmixMosaic). The final version of the code will be archived on Dryad upon acceptance of the manuscript.

## References

Adam AA, Thomas L, Underwood J, Gilmour J, Richards ZT (2022) Population connectivity and genetic offset in the spawning coral *Acropora digitifera* in western australia. Molecular Ecology, 31, 3533–3547.

Aguirre-Liguori JA, Ramírez-Barahona S, Tiffin P, Eguiarte LE (2019) Climate change is predicted to disrupt patterns of local adaptation in wild and cultivated maize. Proceedings of the Royal Society B: Biological Sciences, 286, 20190486.

Ahrens CW, Rymer PD, Miller AD (2026) Genetic offset and vulnerability modeling under climate change scenarios: common misinterpretations and violations of evolutionary principles. Evolution, 80, 15–27.

Aitken SN, Yeaman S, Holliday JA, Wang T, Curtis-McLane S (2008) Adaptation, migration or extirpation: climate change outcomes for tree populations. Evolutionary Applications, 1, 95–111.

Angert AL, Bontrager MG, °Agren J (2020) What do we really know about adaptation at range edges? Annual Review of Ecology, Evolution, and Systematics, 51, 341–361.

Archambeau J, Benito Garzón M, de Miguel M, et al. (2026) Evaluating genomic offset predictions in a forest tree with high population genetic structure. The American Naturalist, 207, 389–414.

Bay RA, Harrigan RJ, Underwood VL, Gibbs HL, Smith TB, Ruegg K (2018) Genomic signals of selection predict climate-driven population declines in a migratory bird. Science, 359, 83–86.

Blanchet FG, Legendre P, Borcard D (2008) Forward selection of explanatory variables. Ecology, 89, 2623–2632.

Blüthgen N, Dicks LV, Forister ML, Outhwaite CL, Slade EM (2023) Insect declines in the anthropocene. Nature Reviews Earth & Environment, 4, 683–686.

Bonnier J, Saéz Laguna E, Françisco T, et al. (2025) Wet season environments drive local adaptation in the timber tree *Dicorynia guianensis* in french guiana. Molecular Ecology, 34, e17759.

Booker TR (2024) The structure of the environment influences the patterns and genetics of local adaptation. Evolution Letters, 8, 787–798.

Borrell JS, Zohren J, Nichols RA, Buggs RJ (2020) Genomic assessment of local adaptation in dwarf birch to inform assisted gene flow. Evolutionary Applications, 13, 161–175.

Bourret A, Leung C, Puncher GN, et al. (2024) Diving into broad-scale and high-resolution population genomics to decipher drivers of structure and climatic vulnerability in a marine invertebrate. Molecular Ecology, 33, e17448.

Bradshaw WE, Holzapfel CM (2006) Evolutionary response to rapid climate change. Science, 312, 1477–1478.

Brady SP, Bolnick DI, Angert AL, et al. (2019) Causes of maladaptation. Evolutionary Applications, 12, 1229–1242.

Bush A, Mokany K, Catullo R, et al. (2016) Incorporating evolutionary adaptation in species distribution modelling reduces projected vulnerability to climate change. Ecology letters, 19, 1468–1478.

Candido-Ribeiro R, Lind BM, Singh P, et al. (2026) Polygenic and redundant architectures of climate-adaptive traits may complicate genomic predictions of maladaptation. *bioRxiv*, pp. 2026–02.

Capblancq T, Fitzpatrick MC, Bay RA, Exposito-Alonso M, Keller SR (2020a) Genomic prediction of (mal) adaptation across current and future climatic landscapes. Annual Review of Ecology, Evolution, and Systematics, 51, 245–269.

Capblancq T, Forester BR (2021) Redundancy analysis: A swiss army knife for landscape genomics. Methods in Ecology and Evolution, 12, 2298–2309.

Capblancq T, Luu K, Blum MG, Bazin E (2018) Evaluation of redundancy analysis to identify signatures of local adaptation. Molecular ecology resources, 18, 1223–1233.

Capblancq T, Morin X, Gueguen M, Renaud J, Lobreaux S, Bazin E (2020b) Climate-associated genetic variation in *Fagus sylvatica* and potential responses to climate change in the french alps. Journal of Evolutionary Biology, 33, 783–796.

Caye K, Jumentier B, Lepeule J, François O (2019) Lfmm 2: fast and accurate inference of gene-environment associations in genome-wide studies. Molecular Biology and Evolution, 36, 852–860.

Chaturvedi S, Lucas LK, Buerkle CA, et al. (2020) Recent hybrids recapitulate ancient hybrid outcomes. Nature communications, 11, 2179.

Chen Y, Chen Y, Shi C, et al. (2018) Soapnuke: a mapreduce acceleration-supported software for integrated quality control and preprocessing of high-throughput sequencing data. Gigascience, 7, gix120.

Chen Y, Jiang Z, Fan P, et al. (2022) The combination of genomic offset and niche modelling provides insights into climate change-driven vulnerability. Nature Communications, 13, 4821.

Dauphin B, Rellstab C, Schmid M, et al. (2021) Genomic vulnerability to rapid climate warming in a tree species with a long generation time. Global change biology, 27, 1181–1195.

Ebbert MT, Wadsworth ME, Staley LA, et al. (2016) Evaluating the necessity of pcr duplicate removal from next-generation sequencing data and a comparison of approaches. BMC bioinformatics, 17, 239.

Edwards CB, Zipkin EF, Henry EH, et al. (2025) Rapid butterfly declines across the united states during the 21st century. Science, 387, 1090–1094.

Exposito-Alonso M, Burbano HA, Bossdorf O, Nielsen R, Weigel D (2019) Natural selection on the *Arabidopsis thaliana* genome in present and future climates. Nature, 573, 126–129.

Fitzpatrick MC, Blois JL, Williams JW, Nieto-Lugilde D, Maguire KC, Lorenz DJ (2018) How will climate novelty influence ecological forecasts? using the quaternary to assess future reliability. Global Change Biology, 24, 3575–3586.

Fitzpatrick MC, Chhatre VE, Soolanayakanahally RY, Keller SR (2021) Experimental support for genomic prediction of climate maladaptation using the machine learning approach gradient forests. Molecular Ecology Resources, 21, 2749–2765.

Fitzpatrick MC, Keller SR (2015) Ecological genomics meets community-level modelling of biodiversity: Mapping the genomic landscape of current and future environmental adaptation. Ecology letters, 18, 1–16.

Fitzpatrick MC, Keller SR, Lotterhos KE (2026) The challenge of genomic forecasting in an era of global change. The American Naturalist, 207, 347–355.

Fordyce J, Nice C, Forister M, Shapiro A (2002) The significance of wing pattern diversity in the lycaenidae: mate discrimination by two recently diverged species. Journal of Evolutionary biology, 15, 871–879.

Forester BR, Lasky JR, Wagner HH, Urban DL (2018) Comparing methods for detecting multilocus adaptation with multivariate genotype–environment associations. Molecular Ecology, 27, 2215–2233.

Forister M, Halsch C, Nice C, et al. (2021) Fewer butterflies seen by community scientists across the warming and drying landscapes of the american west. Science, 371, 1042–1045.

Forister ML, Fordyce JA, Nice CC, Gompert Z, Shapiro AM (2006) Egg morphology varies among populations and habitats along a suture zone in the *Lycaeides idas-melissa* species complex (lepidoptera: Lycaenidae). Annals of the Entomological Society of America, 99, 933–937.

Forister ML, Gompert Z, Fordyce JA, Nice CC (2011) After 60 years, an answer to the question: what is the karner blue butterfly? Biology Letters, 7, 399–402.

Forister ML, Nice CC, Fordyce JA, Gompert Z (2009) Host range evolution is not driven by the optimization of larval performance: the case of lycaeides melissa (lepidoptera: Lycaenidae) and the colonization of alfalfa. Oecologia, 160, 551–561.

Françisco T, Mayol M, Vajana E, et al. (2025) Genomic signatures of climate-driven (mal) adaptation in an iconic conifer, the english yew (*Taxus baccata L*.). Evolutionary Applications, 18, e70160.

François O, Martins H, Caye K, Schoville SD (2016) Controlling false discoveries in genome scans for selection. Molecular Ecology, 25, 454–469.

Frichot E, Schoville SD, Bouchard G, Francois O (2013) Testing for associations between loci and environmental gradients using latent factor mixed models. Molecular Biology and Evolution, 30, 1687–1699.

Gain C, Rhoné B, Cubry P, et al. (2023) A quantitative theory for genomic offset statistics. Molecular Biology and Evolution, 40, msad140.

Garnas JR (2018) Rapid evolution of insects to global environmental change: conceptual issues and empirical gaps. Current Opinion in Insect Science, 29, 93–101.

Gautier M (2015) Genome-wide scan for adaptive divergence and association with population-specific covariates. Genetics, 201, 1555–1579.

Gompert Z, Lucas LK, Fordyce JA, Forister ML, Nice CC (2010) Secondary contact between *Lycaeides idas* and *L. melissa* in the rocky mountains: extensive admixture and a patchy hybrid zone. Molecular Ecology, 19, 3171–3192.

Gompert Z, Lucas LK, Nice CC, Buerkle CA (2013a) Genome divergence and the genetic architecture of barriers to gene flow between *Lycaeides idas* and *L. melissa*. Evolution, 67, 2498–2514.

Gompert Z, Lucas LK, Nice CC, Fordyce JA, Alex Buerkle C, Forister ML (2013b) Geographically multifarious phenotypic divergence during speciation. Ecology and Evolution, 3, 595–613.

Gompert Z, Lucas LK, Nice CC, Fordyce JA, Forister ML, Buerkle CA (2012) Genomic regions with a history of divergent selection affect fitness of hybrids between two butterfly species. Evolution, 66, 2167–2181.

Gompert Z, Nice CC, Fordyce JA, Forister ML, Shapiro AM (2006) Identifying units for conservation using molecular systematics: the cautionary tale of the karner blue butterfly. Molecular Ecology, 15, 1759–1768.

Goodwin KB, Chaturvedi S, Lucas LK, Gompert Z (2026) Genomic forecasts of maladptation in lycaeides butterflies. bioRxiv, pp. 2026–05.

Gougherty AV, Keller SR, Fitzpatrick MC (2021) Maladaptation, migration and extirpation fuel climate change risk in a forest tree species. Nature Climate Change, 11, 166–171.

Gugger PF, Liang CT, Sork VL, Hodgskiss P, Wright JW (2018) Applying landscape genomic tools to forest management and restoration of hawaiian koa (acacia koa) in a changing environment. Evolutionary Applications, 11, 231–242.

Hargreaves AL, Samis KE, Eckert CG (2014) Are species’ range limits simply niche limits writ large? a review of transplant experiments beyond the range. The American Naturalist, 183, 157–173.

Henry EH, Edwards CB, Shirey V, et al. (2025) Twenty years (2000–2020) of butterfly monitoring data across the contiguous united states. Scientific Data, 12, 1869.

Hoffmann AA, Sgró CM (2011) Climate change and evolutionary adaptation. Nature, 470, 479–485.

Hoste A, Capblancq T, Broquet T, et al. (2024) Projection of current and future distribution of adaptive genetic units in an alpine ungulate. Heredity, 132, 54–66.

Ingvarsson PK, Bernhardsson C (2020) Genome-wide signatures of environmental adaptation in european aspen (populus tremula) under current and future climate conditions. Evolutionary applications, 13, 132–142.

IPCC (2014) Climate Change 2014: The Physical Science Basis. Cambridge University Press, Cambridge, United Kingdom and New York, NY, USA.

Jia KH, Zhao W, Maier PA, et al. (2020) Landscape genomics predicts climate change-related genetic offset for the widespread *Platycladus orientalis* (cupressaceae). Evolutionary Applications, 13, 665–676.

Lachmuth S, Capblancq T, Keller SR, Fitzpatrick MC (2023) Assessing uncertainty in genomic offset forecasts from landscape genomic models (and implications for restoration and assisted migration). Frontiers in Ecology and Evolution, 11, 1155783.

Lachmuth S, Capblancq T, Prakash A, Keller SR, Fitzpatrick MC (2024) Novel genomic offset metrics integrate local adaptation into habitat suitability forecasts and inform assisted migration. Ecological Monographs, 94, e1593.

Láruson ÁJ, Fitzpatrick MC, Keller SR, Haller BC, Lotterhos KE (2022) Seeing the forest for the trees: Assessing genetic offset predictions from gradient forest. Evolutionary Applications, 15, 403–416.

Láruson ÁJ, Yeaman S, Lotterhos KE (2020) The importance of genetic redundancy in evolution. Trends in ecology & evolution, 35, 809–822.

Lee-Yaw JA, Kharouba HM, Bontrager M, et al. (2016) A synthesis of transplant experiments and ecological niche models suggests that range limits are often niche limits. Ecology letters, 19, 710–722.

Li H (2011) A statistical framework for snp calling, mutation discovery, association mapping and population genetical parameter estimation from sequencing data. Bioinformatics, 27, 2987–2993.

Li H, Durbin R (2009) Fast and accurate short read alignment with burrows–wheeler transform. Bioinformatics, 25, 1754–1760.

Li H, Handsaker B, Wysoker A, et al. (2009) The sequence alignment/map format and samtools. Bioinformatics, 25, 2078–2079.

Lind BM, Candido-Ribeiro R, Singh P, et al. (2024) How useful is genomic data for predicting maladaptation to future climate? Global Change Biology, 30, e17227.

Lind BM, Lotterhos KE (2025) The accuracy of predicting maladaptation to new environments with genomic data. Molecular Ecology Resources, 25, e14008.

Lotterhos KE (2023) The paradox of adaptive trait clines with nonclinal patterns in the underlying genes. Proceedings of the National Academy of Sciences, 120, e2220313120.

Lotterhos KE (2024) Principles in experimental design for evaluating genomic forecasts. Methods in Ecology and Evolution, 15, 1466–1482.

Lucas L, Fordyce J, Nice C (2008) Patterns of genitalic morphology around suture zones in north american lycaeides (lepidoptera: Lycaenidae): implications for taxonomy and historical biogeography. Annals of the Entomological Society of America, 101, 172–180.

Lucas LK, Nice CC, Gompert Z (2018) Genetic constraints on wing pattern variation in lycaeides butterflies: A case study on mapping complex, multifaceted traits in structured populations. Molecular Ecology Resources, 18, 892–907.

Marchi M, Bucci G, Iovieno P, Ray D (2024) Climatedt: a global scale-free dynamic downscaling portal for historic and future climate data. Environments, 11, 82.

Martins K, Gugger PF, Llanderal-Mendoza J, et al. (2018) Landscape genomics provides evidence of climate-associated genetic variation in mexican populations of *Quercus rugosa*. Evolutionary Applications, 11, 1842–1858.

Mathieson I, McVean G (2013) Estimating selection coefficients in spatially structured populations from time series data of allele frequencies. Genetics, 193, 973–984.

McKenna A, Hanna M, Banks E, et al. (2010) The genome analysis toolkit: a mapreduce framework for analyzing next-generation dna sequencing data. Genome research, 20, 1297–1303.

McLennan EA, Kovacs TG, Silver LW, et al. (2025) Genomics identifies koala populations at risk across eastern australia. Ecological Applications, 35, e3062.

Morán-Ordóñez A, Briscoe NJ, Wintle BA (2018) Modelling species responses to extreme weather provides new insights into constraints on range and likely climate change impacts for australian mammals. Ecography, 41, 308–320.

Nabokov V (1949) The nearctic members of lycaeides hübner (lycaenidae, lepidoptera). Bulletin of the Museum of Comparative Zoology, 101, 479–541.

Nadeau CP, Urban MC (2019) Eco-evolution on the edge during climate change. Ecography, 42, 1280–1297.

Nadeau CP, Urban MC, Bridle JR (2017) Climates past, present, and yet-to-come shape climate change vulnerabilities. Trends in Ecology & Evolution, 32, 786–800.

Nice C, Shapiro A (1999) Molecular and morphological divergence in the butterfly genus lycaeides (lepidoptera: Lycaenidae) in north america: evidence of recent speciation. Journal of Evolutionary Biology, 12, 936–950.

Nice CC, Forister ML, Harrison JG, et al. (2019) Extreme heterogeneity of population response to climatic variation and the limits of prediction. Global Change Biology, 25, 2127–2136.

Nice CC, Gompert Z, Fordyce JA, Forister ML, Lucas LK, Buerkle CA (2013) Hybrid speciation and independent evolution in lineages of alpine butterflies. Evolution, 67, 1055–1068.

Rêgo A, Baur J, Girard-Tercieux C, de la Paz Celorio-Mancera M, Stelkens R, Berger D (2025) Repeatability of evolution and genomic predictions of temperature adaptation in seed beetles. Nature Ecology & Evolution, 9, 1061–1074.

Reis GA, Forister ML, Halsch CA, Dittemore CM, Shapiro AM, Gompert Z (2026) Climate and species traits give rise to complex phenological dynamics. Ecology, 107, e70297.

Rellstab C, Dauphin B, Exposito-Alonso M (2021) Prospects and limitations of genomic offset in conservation management. Evolutionary Applications, 14, 1202–1212.

Rellstab C, Gugerli F, Eckert AJ, Hancock AM, Holderegger R (2015) A practical guide to environmental association analysis in landscape genomics. Molecular Ecology, 24, 4348–4370.

Rellstab C, Zoller S, Walthert L, et al. (2016) Signatures of local adaptation in candidate genes of oaks (*Quercus spp.*) with respect to present and future climatic conditions. Molecular Ecology, 25, 5907–5924.

Rhoné B, Defrance D, Berthouly-Salazar C, et al. (2020) Pearl millet genomic vulnerability to climate change in west africa highlights the need for regional collaboration. Nature Communications, 11, 5274.

Rudman SM, Greenblum SI, Rajpurohit S, et al. (2022) Direct observation of adaptive tracking on ecological time scales in drosophila. Science, 375, eabj7484.

Ruegg K, Bay RA, Anderson EC, et al. (2018) Ecological genomics predicts climate vulnerability in an endangered southwestern songbird. Ecology Letters, 21, 1085–1096.

Sang Y, Long Z, Dan X, et al. (2022) Genomic insights into local adaptation and future climate-induced vulnerability of a keystone forest tree in east asia. Nature Communications, 13, 6541.

Simon A, Coop G (2024) The contribution of gene flow, selection, and genetic drift to five thousand years of human allele frequency change. Proceedings of the National Academy of Sciences, 121, e2312377121.

Simon JC, Peccoud J (2018) Rapid evolution of aphid pests in agricultural environments. Current opinion in insect science, 26, 17–24.

Smith TB, Fuller TL, Zhen Y, et al. (2021) Genomic vulnerability and socio-economic threats under climate change in an african rainforest bird. Evolutionary Applications, 14, 1239–1247.

Talavera G, Lukhtanov VA, Pierce NE, Vila R (2013) Establishing criteria for higher-level classification using molecular data: the systematics of polyommatus blue butterflies (lepidoptera, lycaenidae). Cladistics, 29, 166–192.

Taron D, Ries L (2015) Butterfly monitoring for conservation. In: Butterfly Conservation in North America: Efforts to Help Save Our Charismatic Microfauna (ed. Daniels JC), pp. 35–57, Springer Netherlands, Dordrecht.

Theraroz A, Guadaño-Peyrot C, Archambeau J, et al. (2024) The genetic consequences of population marginality: A case study in maritime pine. Diversity and Distributions, 30, e13910.

Trujillo L, Banse P, Beslon G (2022) Getting higher on rugged landscapes: Inversion mutations open access to fitter adaptive peaks in NK fitness landscapes. PLoS Computational Biology, 18, e1010647.

Urban MC (2015) Accelerating extinction risk from climate change. Science, 348, 571–573.

Van Deurs S, Reutimann O, Luqman H, et al. (2025) Genomic signatures of adaptation across a precipitation gradient from niche centre to niche edge. Molecular Ecology, 34, e17696.

Vanhove M, Pina-Martins F, Coelho AC, et al. (2021) Using gradient forest to predict climate response and adaptation in cork oak. Journal of Evolutionary Biology, 34, 910–923.

Vasimuddin M, Misra S, Li H, Aluru S (2019) Efficient architecture-aware acceleration of bwa-mem for multicore systems. In: 2019 IEEE international parallel and distributed processing symposium (IPDPS), pp. 314–324, IEEE.

Verrico BM, Capblancq T, Fitzpatrick MC, Keller SR (2026) Reciprocal evaluation of genomic offset predictions of climate maladaptation with independent empirical datasets. The American Naturalist, 207, 415–432.

Wagner DL, Grames EM, Forister ML, Berenbaum MR, Stopak D (2021) Insect decline in the anthropocene: Death by a thousand cuts. Proceedings of the National Academy of Sciences, 118, e2023989118.

Wang T, Hamann A, Spittlehouse D, Carroll C (2016) Locally downscaled and spatially customizable climate data for historical and future periods for north america. PloS one, 11, e0156720.

Wu N, Xiao Q, Liao Z, et al. (2025) Landscape genomics provides insights into climate change-driven vulnerability in torrent frogs (ranidae: Amolops). Molecular Ecology, 34, e17807.

Yeaman S (2015) Local adaptation by alleles of small effect. The American Naturalist, 186, S74–S89.

