## Supplementary Materials for "Genomic offset is not predictive of recent demographic trends in *Lycaeides* butterflies"

### Supplementary methods

#### Library preparation, sequencing and SNP filtering

Sequencing was performed by BGI using DNB-based sequencing technology (Drmanac *et al.*, 2010; Li *et al.*, 2019). Briefly, DNA was fragmented using a Covaris ultrasonicator, and fragment size distributions were assessed with an Agilent 2100 Bioanalyzer. Fragmented DNA underwent end repair and A-tailing, followed by magnetic bead purification. Indexed sequencing adapters were ligated to A-tailed fragments, and excess adapters were removed by bead-based purification. Libraries were amplified using limited-cycle PCR with KAPA HiFi HotStart DNA Polymerase. Amplified libraries were circularized following the MGI protocol and subsequently amplified into DNA nanoballs (DNBs) via rolling-circle amplification. Sequencing was conducted on the DNBseq platform using paired-end 100 bp reads, targeting approximately 100× genome coverage per library. Raw reads were filtered using SOAPnuke to remove adapter contamination, low-quality reads, and reads containing excessive ambiguous bases (Chen *et al.*, 2018). Reads were discarded if >25% of bases matched adapter sequences, ≥40% of bases had a Phred quality score <10, or ambiguous nucleotides (Ns) exceeded 0.1% of the read.
Filtered reads were aligned to the *L. melissa* reference genome using bwa-mem2 (v2.0pre2) with default parameters (Li & Durbin, 2009; Vasimuddin *et al.*, 2019). PCR duplicates were removed using the collate, fixmate, and markdup commands in SAMtools (v1.16) (Ebbert *et al.*, 2016; Li *et al.*, 2009). SNPs were called using the bcftools consensus caller (-c; v1.16), excluding insertion–deletion polymorphisms, bases with quality scores <30, and alignments with mapping quality <20 (Li, 2011). SNPs were retained only if the probability that all populations were fixed for the reference allele was <1%. Additional

filtering was performed using GATK (v4.1.4.1), excluding non-biallelic SNPs, SNPs with total read depth <1,450, and SNPs with mapping quality <30 (McKenna *et al.*, 2010).

#### Climate-associated SNPs

Climate-associated SNPs were identified using four common GEA methods; LFMM, BayPass, RDA, and partial RDA (pRDA) which differ in statistical framework (Bayesian vs. frequentist), dimensionality (univariate vs. multivariate), and whether they correct for population structure (Table S2). Baseline climate data (1961–1990), assumed to represent conditions to which populations were locally adapted, were used as predictor variables in all analyses.

Population structure was characterized using principal component analysis (PCA) on centered and scaled SNP allele frequencies. The first three principal components were retained, explaining 35.1% of the total genetic variance (PC1: 15.52%, PC2: 11.25%, PC3: 8.32%). This choice was supported by the scree plot (Figure S1a), which showed a clear decline in explained variance beyond the third component, indicating diminishing returns from additional axes. Retaining three components captures the major axes of population structure while minimizing overfitting and loss of power.

To assess whether spatial autocorrelation required explicit correction beyond population structure, geographic structure was quantified using distance-based Moran's Eigenvector Maps (dbMEM) implemented in the adespatial package in R. The procedure returned a single eigenvector (MEM1), reflecting a single broad-scale spatial gradient across sampling sites. When modelled alone, MEM1 explained a significant proportion of allele frequency variation ($R^2 = 0.149$, $p = 0.001$). However, its contribution was substantially reduced after conditioning on population structure (PC1–PC3) and climate variables (BIO10, BIO13, BIO15), explaining only a small and marginally significant fraction of variation ($R^2 = 0.024$, $p = 0.05$; Table S3). This reduction indicates that spatial structure is largely captured by population structure already included in the models. Accordingly, explicit correction for spatial autocorrelation was not considered necessary, and population structure alone was used to account for non-independence among populations. This is consistent with Rellstab *et al.* (2015), who caution that jointly accounting for both population structure and spatial autocorrelation may be overly conservative and reduce power to detect true genotype–environment associations.

##### Redundancy Analysis (RDA)

RDA, a multivariate constrained ordination method, was used to model associations between SNP allele frequencies (response variables) and climatic variables (predictor variables). Analyses followed procedures described in Capblancq *et al.* (2018) and Capblancq & Forester (2021) and were implemented using the vegan package in R. Climate-associated SNPs were identified based on locus extremeness in RDA space, quantified using Mahalanobis distances from the centroid of the first two RDA axes. These two axes represented the primary gradients of climate-constrained genetic variation, capturing a cumulative 34.2% of the constrained variance (RDA1: 23.2%; RDA2: 11.0%) (Figure S1b). P-values were computed using the rdadapt package (Capblancq *et al.*, 2024) and adjusted for multiple testing using a false discovery rate (FDR) of 5%. SNPs identified by RDA were hereafter referred to as RDA SNPs.

##### Partial Redundancy Analysis (pRDA)

Partial RDA was implemented identically to RDA, with the exception that population structure was accounted for by including the first three principal components as conditioning variables. SNPs identified by pRDA were hereafter referred to as pRDA SNPs.

##### Latent Factor Mixed Models (LFMM)

LFMMs were used as a univariate approach to test for associations between SNP allele frequencies and climatic variables while accounting for unobserved confounding effects of population structure via latent factors. LFMM analyses were conducted using ridge regression implemented in the lfmm R package following Caye *et al.* (2019). Three latent factors were included to account for population structure, consistent with the pRDA approach. P-values were calibrated using the genomic inflation factor (GIF) and adjusted for multiple testing using an FDR of 5%. SNPs identified by LFMM were hereafter referred to as LFMM SNPs.

##### BayPass

BayPass v3.1 a Bayesian univariate GEA method was used to identify climate-associated SNPs following Gautier (2015). Population structure was first estimated using the core model (no covariates) to infer the covariance matrix of allele frequencies among populations. The standard covariate model was then used to test for associations between SNP allele frequencies and climatic variables while accounting for this covariance structure. Bayes factors were estimated using an importance sampling approximation (Coop *et al.*, 2010). Three independent runs were performed using different random seeds to ensure convergence, and median Bayes factors across runs were computed as recommended (Gautier, 2015; Gautier *et al.*, 2018). SNPs with a median Bayes factor >20 deciban units for association with any climatic variable, indicative of decisive evidence under Jeffreys' scale (Jeffreys, 1961), were classified as climate-associated. SNPs identified by BayPass were hereafter referred to as BayPass SNPs.

#### Estimation of population trends

We estimated population trends of *Lycaeides* butterfly populations at North American Butterfly Association (NABA) monitoring sites and extended these estimates to populations sampled for genomic analysis. Trends at monitoring sites were quantified directly using the NABA dataset (Figure 1c, 4b). Trends at populations sampled for genomic analysis were subsequently derived through spatial interpolation from nearby monitoring locations. The modeling framework for each dataset and the interpolation procedure are described below.

#### North American Butterfly Association (NABA) dataset

This NABA dataset encompasses hundreds of survey circles across the United States and Canada. Methods regarding surveys are described extensively in Edwards *et al.* (2025), Forister *et al.* (2021), and Henry *et al.* (2025). Briefly, volunteers survey 15 mile (24.14km) diameter circles during the summer at different locations, identifying butterflies and recording the number of individuals of each species observed. Total time spent surveying at each location is also noted. Here, we used data collected in the western United States for all *Lycaeides* spp. At sites where multiple nominal species were counted, we summed counts of all nominal species together. We restricted our analyses to sites that have been monitored for at least ten years and where at least five individuals of any *Lycaeides* butterfly have been counted in five years, to exclude sites where these species are strays (Figure 1c, 4b). To estimate population trends of *Lycaeides* at each site, we used a hierarchical Bayesian framework. Counts of individuals at site $i$ in year $j$ ($C_{ij}$) were modeled using a Poisson generalized linear mixed model. Year, survey effort (hours), and survey date, all z-standardized, were included as predictors in a design matrix $\mathbf{X}_j$ (Equation 1, 2). Our hierarchical framework allows both intercepts and slopes to vary among sites, capturing site-specific effects of each predictor variable. Here, $\alpha_i$ represents the site-specific intercept, and $\boldsymbol{\beta}_i$ represents the vector of site-specific effects for the predictor variables at site $i$. In particular, the site-specific effect of year in $\boldsymbol{\beta}_i$ quantifies the temporal trend in *Lycaeides* abundance at site $i$. Site-specific intercepts and predictor effects were modeled hierarchically as arising from normal distributions, with their means and variances governed by species-level hyperparameters assigned weakly informative priors (Equations 3, 4, 5, 6, 7, 8).

$$C_{ij} \sim \text{Poisson}(\lambda_{ij}) \tag{1}$$

$$\log(\lambda_{ij}) = \alpha_i + \boldsymbol{\beta}_i^\top \mathbf{X}_j \tag{2}$$

$$\alpha_i \sim \text{Normal}(\mu_\alpha, \sigma_\alpha) \tag{3}$$

$$\boldsymbol{\beta}_i \sim \text{MVN}(\boldsymbol{\mu}_\beta, \boldsymbol{\Sigma}_\beta) \tag{4}$$

$$\mu_\alpha \sim \text{Normal}(0, 20) \tag{5}$$

$$\sigma_\alpha \sim \text{Gamma}(2, 0.1) \tag{6}$$

$$\boldsymbol{\mu}_\beta \sim \text{Normal}(0, 20) \tag{7}$$

$$\boldsymbol{\Sigma}_\beta \sim \text{Gamma}(2, 0.1) \tag{8}$$

For modeling the NABA dataset, we sampled posteriors using Hamiltonian Monte Carlo in Stan using the Rstan package. We ran four chains with 20,000 iterations and the first 10,000 as burn-in. We assessed the convergence of models using Gelman-Rubin diagnostics and trace plots. Posterior distributions for site-specific estimates were summarized by estimating their medians. The effect of year in the model determined the population trend of each site, with a negative effect depicting a decline in abundance and a positive effect depicting an increase in abundance (Figure 4b).

**Genomic dataset**

Demographic trends were estimated at populations sampled for genomic analysis using inverse distance weighting (IDW) based on trend estimates from NABA monitoring sites (Bay *et al.*, 2018; Ruegg *et al.*, 2018). To ensure interpolation was restricted to spatially relevant areas, we calculated geodesic distances between each genomic sampled population and all NABA sites using the Vincenty ellipsoid method implemented in the geosphere R package. Populations sampled for genomic analysis located more than 50 km from any NABA site were excluded from IDW interpolation. IDW interpolation was conducted using the gstat R package with a weighting power parameter of 2 (inverse squared distance). To assess sensitivity to the number of neighbors, interpolations were performed using 2 to 10 nearest NABA sites. Correlations among predicted trends across neighbor counts were consistently high ($r = 1.0$). We therefore used five nearest neighbors for the final interpolation.

### Supplementary Tables

Table S1: Site locations and sample sizes of populations sampled for genomic analysis

| ID | Longitude | Latitude | Sample Size |
|---|---|---|---|
| ABM | -111.6230 | 40.58700 | 48 |
| BCR | -110.5530 | 43.30100 | 48 |
| BHP | -118.3910 | 37.35800 | 48 |
| BKM | -120.2900 | 41.69000 | 33 |
| BLD | -109.7161 | 43.52245 | 50 |
| BNP | -110.7210 | 44.93400 | 49 |
| BTB | -110.6820 | 43.63800 | 48 |
| CLH | -118.1900 | 37.46000 | 36 |
| CP | -120.0200 | 38.71000 | 48 |
| EP | -120.2200 | 41.26000 | 48 |
| GNP | -111.2200 | 45.43000 | 56 |
| HJ | -120.0734 | 39.77972 | 48 |
| HNV | -110.4950 | 44.68200 | 48 |
| HUM | -107.7466 | 44.74925 | 63 |
| LAE | -119.4800 | 38.28000 | 48 |
| LS | -120.2400 | 38.63000 | 48 |
| MR | -119.9300 | 39.32000 | 48 |
| MTU | -122.3770 | 41.77200 | 48 |
| PSP | -110.8490 | 42.74700 | 48 |
| RNV | -110.8850 | 43.59600 | 32 |
| SHC | -122.1640 | 41.88100 | 46 |
| SIN | -107.0925 | 41.85110 | 46 |
| SKI | -110.9227 | 43.50942 | 48 |
| SUV | -120.1000 | 41.28000 | 59 |
| TBY | -109.4500 | 44.95000 | 24 |
| TIC | -119.2600 | 37.97000 | 48 |
| USL | -110.3328 | 43.58290 | 48 |
| VE | -120.0000 | 39.51000 | 48 |
| YG | -120.6000 | 39.32000 | 48 |

Table S2: Comparison of genotype–environment association methods used to identify climate-associated SNPs, across key methodological properties. Methods differ in whether they account for population structure and whether they are univariate or multivariate in their treatment of genotype–environment associations. Shaded (dark) cells indicate that a method possesses the corresponding property; unshaded (white) cells indicate that it does not.

| **Methods** | **Population structure** | **Univariate** | **Multivariate** |
|---|---|---|---|
| RDA | | | ■ |
| pRDA | ■ | | ■ |
| BAYPASS | ■ | ■ | |
| LFMM | ■ | ■ | |

Table S3: Redundancy analysis (RDA) models evaluating the contribution of spatial autocorrelation to genome-wide allele frequency variation. Spatial structure was quantified using distance-based Moran's Eigenvector Maps (dbMEM); the procedure returned a single eigenvector (MEM1), reflecting a single broad-scale spatial gradient across sampling sites. The marginal model estimates the total variance attributable to MEM1 alone. The partial model isolates the unique contribution of MEM1 after conditioning on population structure (PC1–PC3) and climate (BIO10, BIO13, BIO15), testing whether this spatial gradient explains genetic variation independently of these covariates.

| **Model** | **Variance** | $\mathbf{R^2}$ | **p(>F)** |
|---|---|---|---|
| Spatial autocorrelation only: MEM1 | 27,314 | 0.149 | 0.001 |
| Spatial autocorrelation \| (population structure + climate): MEM1 \| (PC1–PC3 + BIO10 + BIO13 + BIO15) | 4,404 | 0.024 | 0.050 |

### Supplementary Figures

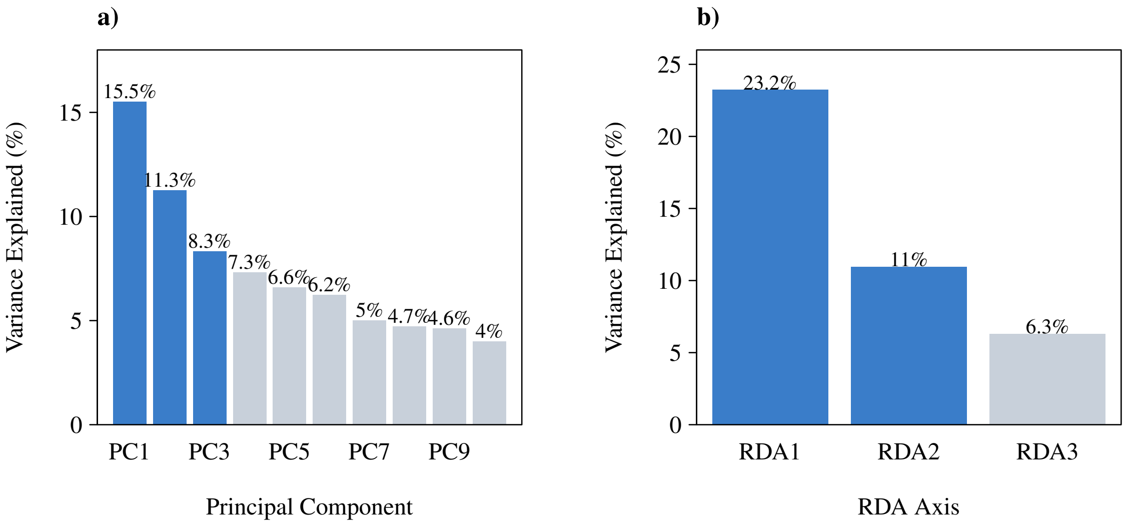


Figure S1: (a) Principal component analysis (PCA) scree plot showing variance explained by the first 10 principal components (PC). Blue bars indicate the three retained components (PC1–PC3), which together explain 35.1% of genetic variation. (b) Variance explained by constrained redundancy analysis (RDA) axes. Blue bars indicate RDA1 and RDA2, which capture a cumulative 34.2% of the constrained variance, representing the primary gradients of climate-associated genetic variation.

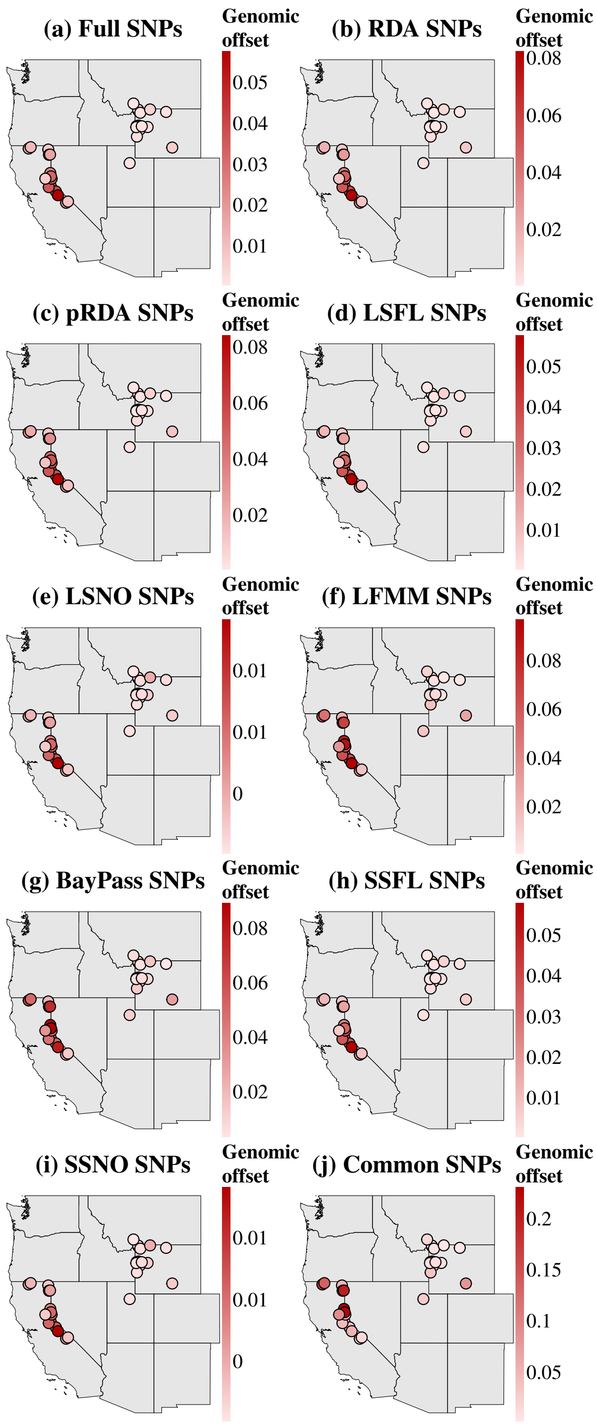


Figure S2: Spatial distribution of weighted environmental distance (WED) estimates across populations sampled for genomic analysis using generalized dissimilarity modeling (GDM). WED was estimated for each of the populations sampled for genomic analysis using ten SNP sets: (a) full SNP dataset (Full SNPs), (b–e) climate-associated SNPs identified by pRDA, RDA, LFMM, and BayPass, (f) SNPs identified by all four GEA methods (Common SNPs), (g–h) large and small random samples of the full dataset (LSFL and SSFL), and (i–j) large and small random subsets of non climate-associated SNPs (LSNO and SSNO). Darker red indicates higher genomic offset.

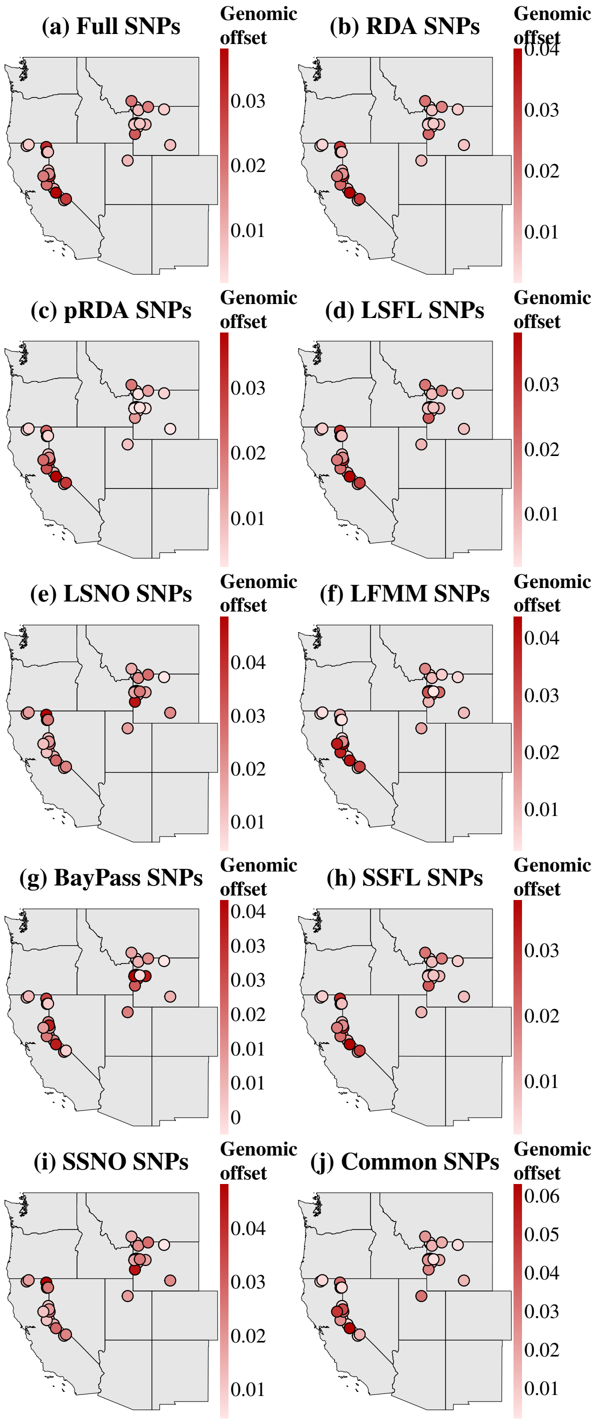


Figure S3: Spatial distribution of weighted environmental distance (WED) estimates across populations sampled for genomic analysis using redundancy analysis (RDA). WED was estimated for each of the populations sampled for genomic analysis using ten SNP sets: (a) full SNP dataset (Full SNPs), (b–e) climate-associated SNPs identified by pRDA, RDA, LFMM, and BayPass, (f) SNPs identified by all four GEA methods (Common SNPs), (g–h) large and small random samples of the full dataset (LSFL and SSFL), and (i–j) large and small random subsets of non climate-associated SNPs (LSNO and SSNO). Darker red indicates higher genomic offset.

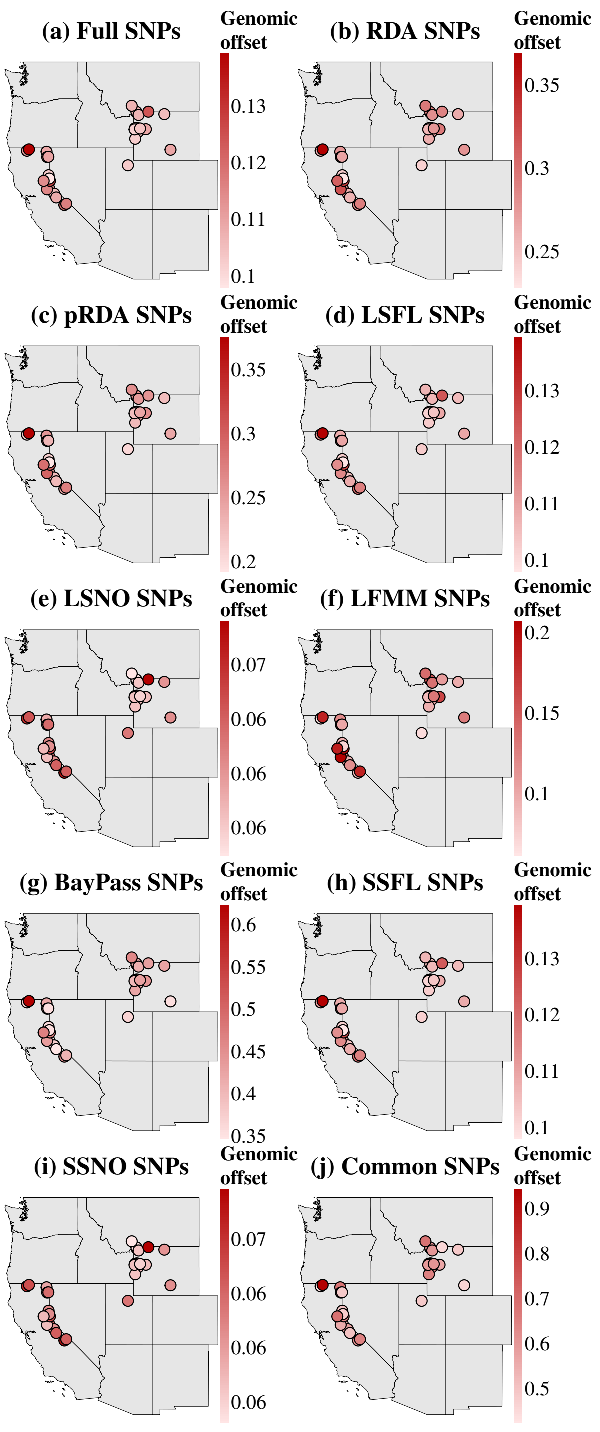


Figure S4: Spatial distribution of current risk of nonadaptedness (C-RONA) estimates across populations sampled for genomic analysis using redundancy analysis (RDA). C-RONA was estimated for each of the populations sampled for genomic analysis using ten SNP sets: (a) full SNP dataset (Full SNPs), (b–e) climate-associated SNPs identified by pRDA, RDA, LFMM, and BayPass, (f) SNPs identified by all four GEA methods (Common SNPs), (g–h) large and small random samples of the full dataset (LSFL and SSFL), and (i–j) large and small random subsets of non climate-associated SNPs (LSNO and SSNO). Darker red indicates higher genomic offset.

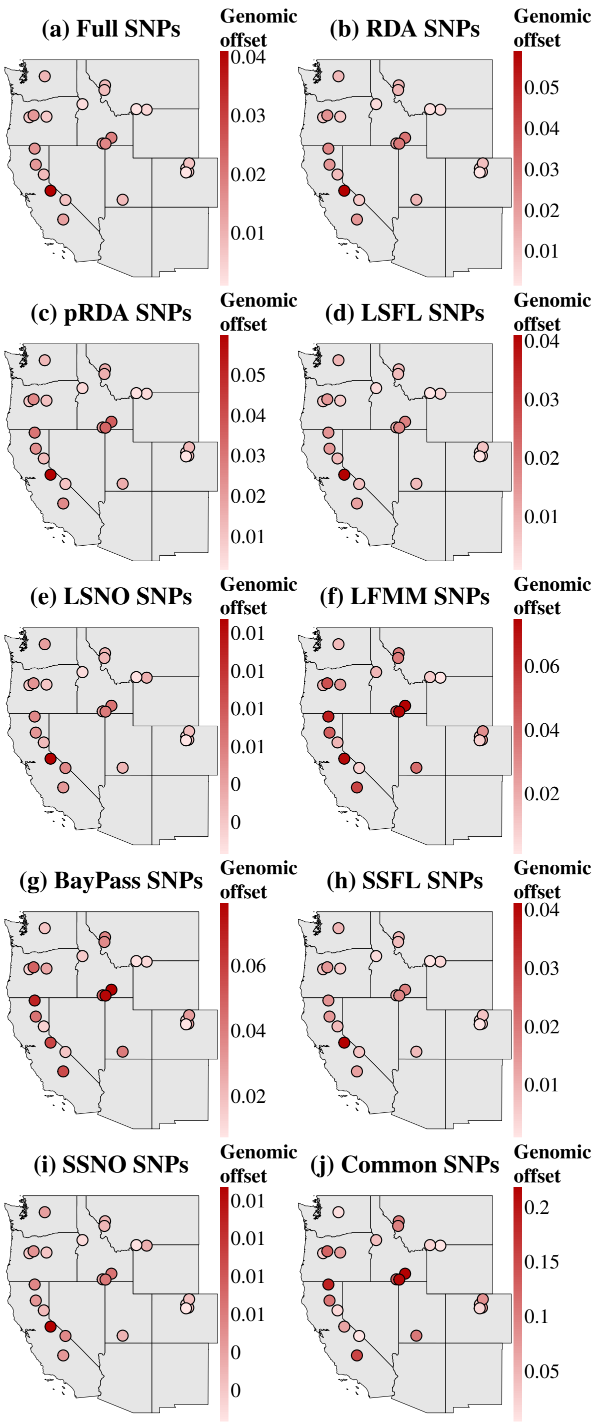


Figure S5: Spatial distribution of weighted environmental distance (WED) estimates across monitored sites using generalized dissimilarity modeling (GDM). WED was extrapolated to North American Butterfly Association (NABA) monitoring sites by applying GEA relationships estimated from populations sampled for genomic analysis. Estimates are shown for ten SNP sets: (a) full SNP dataset (Full SNPs), (b–e) climate-associated SNPs identified by pRDA, RDA, LFMM, and BayPass, (f) SNPs identified by all four GEA methods (Common SNPs), (g–h) large and small random samples of the full dataset (LSFL and SSFL), and (i–j) large and small random subsets of non climate-associated SNPs (LSNO and SSNO). Darker red indicates higher genomic offset.

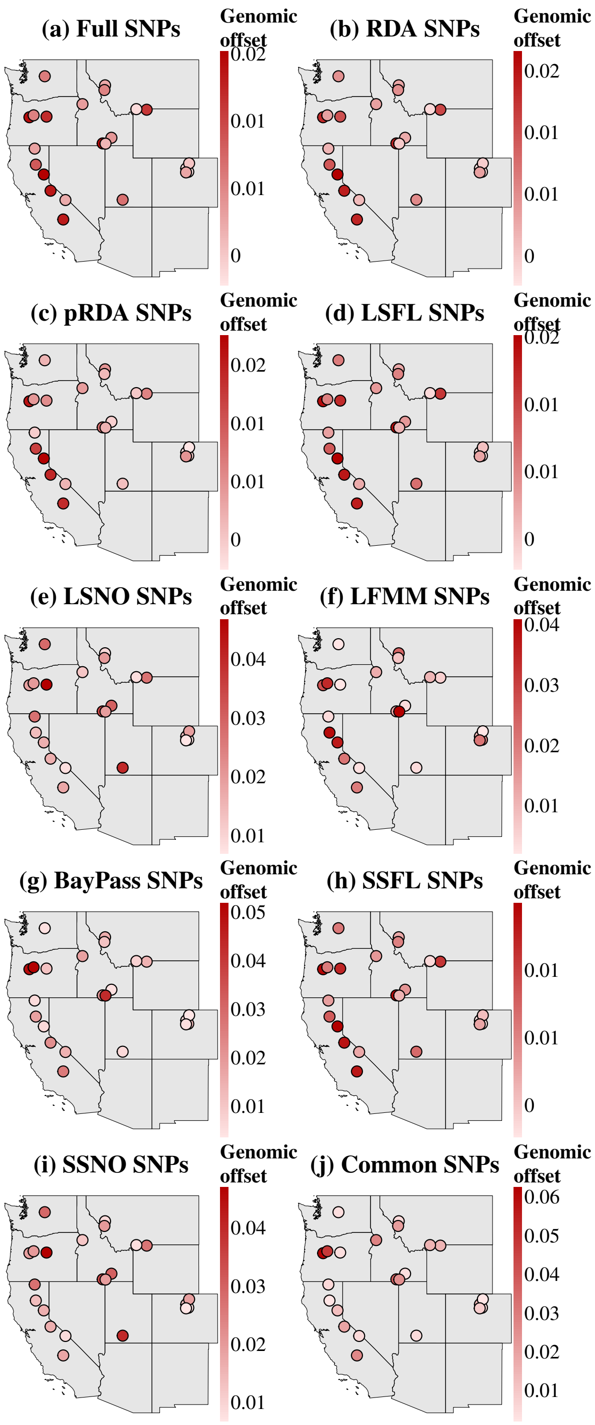


Figure S6: Spatial distribution of weighted environmental distance (WED) estimates across monitored sites using redundancy analysis (RDA). WED was extrapolated to North American Butterfly Association (NABA) monitoring sites by applying GEA relationships estimated from populations sampled for genomic analysis. Estimates are shown for ten SNP sets: (a) full SNP dataset (Full SNPs), (b–e) climate-associated SNPs identified by pRDA, RDA, LFMM, and BayPass, (f) SNPs identified by all four GEA methods (Common SNPs), (g–h) large and small random samples of the full dataset (LSFL and SSFL), and (i–j) large and small random subsets of non climate-associated SNPs (LSNO and SSNO). Darker red indicates higher genomic offset.

### References

Bay RA, Harrigan RJ, Underwood VL, Gibbs HL, Smith TB, Ruegg K (2018) Genomic signals of selection predict climate-driven population declines in a migratory bird. *Science*, **359**, 83–86.

Capblancq T, Forester BR (2021) Redundancy analysis: A swiss army knife for landscape genomics. *Methods in Ecology and Evolution*, **12**, 2298–2309.

Capblancq T, Gueguen M, Forester B (2024) *rdadapt: The multiple uses of RDA in landscape genomics*. R package version 1.0.0.

Capblancq T, Luu K, Blum MG, Bazin E (2018) Evaluation of redundancy analysis to identify signatures of local adaptation. *Molecular ecology resources*, **18**, 1223–1233.

Caye K, Jumentier B, Lepeule J, François O (2019) Lfmm 2: fast and accurate inference of gene-environment associations in genome-wide studies. *Molecular Biology and Evolution*, **36**, 852–860.

Chen Y, Chen Y, Shi C, *et al.* (2018) Soapnuke: a mapreduce acceleration-supported software for integrated quality control and preprocessing of high-throughput sequencing data. *Gigascience*, **7**, gix120.

Coop G, Witonsky D, Di Rienzo A, Pritchard JK (2010) Using environmental correlations to identify loci underlying local adaptation. *Genetics*, **185**, 1411–1423.

Drmanac R, Sparks AB, Callow MJ, *et al.* (2010) Human genome sequencing using unchained base reads on self-assembling dna nanoarrays. *Science*, **327**, 78–81.

Ebbert MT, Wadsworth ME, Staley LA, *et al.* (2016) Evaluating the necessity of pcr duplicate removal from next-generation sequencing data and a comparison of approaches. *BMC bioinformatics*, **17**, 239.

Edwards CB, Zipkin EF, Henry EH, *et al.* (2025) Rapid butterfly declines across the united states during the 21st century. *Science*, **387**, 1090–1094.

Forister M, Halsch C, Nice C, *et al.* (2021) Fewer butterflies seen by community scientists across the warming and drying landscapes of the american west. *Science*, **371**, 1042–1045.

Gautier M (2015) Genome-wide scan for adaptive divergence and association with population-specific covariates. *Genetics*, **201**, 1555–1579.

Gautier M, Yamaguchi J, Foucaud J, *et al.* (2018) The genomic basis of color pattern polymorphism in the harlequin ladybird. *Current Biology*, **28**, 3296–3302.

Henry EH, Edwards CB, Shirey V, *et al.* (2025) Twenty years (2000–2020) of butterfly monitoring data across the contiguous united states. *Scientific Data*, **12**, 1869.

Jeffreys H (1961) *Theory of Probability*. 3rd edn., Oxford University Press, London and New York and Oxford.

Li H (2011) A statistical framework for snp calling, mutation discovery, association mapping and population genetical parameter estimation from sequencing data. *Bioinformatics*, **27**, 2987–2993.

Li H, Durbin R (2009) Fast and accurate short read alignment with burrows–wheeler transform. *Bioinformatics*, **25**, 1754–1760.

Li H, Handsaker B, Wysoker A, *et al.* (2009) The sequence alignment/map format and samtools. *Bioinformatics*, **25**, 2078–2079.

Li Q, Zhao X, Zhang W, *et al.* (2019) Reliable multiplex sequencing with rare index mis-assignment on dnb-based ngs platform. *BMC genomics*, **20**, 215.

McKenna A, Hanna M, Banks E, *et al.* (2010) The genome analysis toolkit: a mapreduce framework for analyzing next-generation dna sequencing data. *Genome research*, **20**, 1297–1303.

Rellstab C, Gugerli F, Eckert AJ, Hancock AM, Holderegger R (2015) A practical guide to environmental association analysis in landscape genomics. *Molecular Ecology*, **24**, 4348–4370.

Ruegg K, Bay RA, Anderson EC, *et al.* (2018) Ecological genomics predicts climate vulnerability in an endangered southwestern songbird. *Ecology Letters*, **21**, 1085–1096.

Vasimuddin M, Misra S, Li H, Aluru S (2019) Efficient architecture-aware acceleration of bwa-mem for multicore systems. In: *2019 IEEE international parallel and distributed processing symposium (IPDPS)*, pp. 314–324, IEEE.
